# Distributed neuronal ensembles support episodic-like memory retrieval

**DOI:** 10.64898/2026.06.15.727203

**Authors:** Stefano Guglielmo, Marco Scantamburlo, Niccolò Di Nardo, Michel C. van den Oever, Raffaele Mazziotti, Tommaso Pizzorusso, Nicola Origlia

**Affiliations:** BIO@SNS Laboratory, Scuola Normale Superiore, Pisa, Italy; Institute of Neuroscience, National Research Council, Pisa, Italy; Department of Molecular and Cellular Neurobiology, Center for Neurogenomics and Cognitive Research (CNCR), Amsterdam Neuroscience, Vrije Universiteit Amsterdam, Amsterdam, the Netherlands; Department of Developmental Neuroscience, IRCCS Stella Maris Foundation, Pisa, Italy; Department of Neuroscience, Psychology, Pharmacology, and Child Health, University of Florence, Florence, Italy

## Abstract

**Memory is proposed to depend on neuronal ensembles distributed across multiple brain regions, yet how episodic-like memories are organized across the brain remains unclear. Here we investigated the brain-wide organization of object-place-context (OPC) memory in mice. By mapping c-Fos activation across the brain during memory recall and comparing it to multiple control conditions, we identified a set of brain regions selectively engaged during episodic-like memory retrieval, consistent with the recruitment of a fronto-posterior medial network. Chemogenetic manipulation of learning-activated neuronal ensembles revealed that the selected brain regions are necessary for memory retrieval. Within this network, the retrosplenial cortex emerged as a key region required for successful memory recall, with neuronal ensembles exhibiting properties of engram cells. Electrophysiological recordings during recall revealed distinct novelty-related dynamics across regions. In mPFC, theta power increased during novel exploration, while firing rate increased both immediately before and during interaction with the novel object configuration. In RSP, novelty was associated with a sharp increase in firing rate during exploration. Together, our findings suggest a distributed organization supporting episodic-like memory retrieval in mice, in which a posterior-medial network is activated and necessary for a successful behavioral expression.**

## Introduction

Memory is a fundamental function of the nervous system, highly conserved across species, that shapes adaptive behavior and supports flexible responses to a changing environment. Memory can be divided into distinct processes such as formation, consolidation, retrieval, and forgetting, which induce changes at multiple levels, from molecular and cellular to neural circuits^1,2^. This enduring physical trace of experience, known as memory engram, is formed by specific populations of neurons called engram cells. These neurons are activated during learning, undergo lasting cellular changes, and their reactivation by a subset of the original learning stimuli results in memory recall^3–5^. While early studies identified engram cells in localized brain regions, growing findings suggest that individual memories are not stored in isolated neuronal populations but are represented by distributed ensembles across multiple brain areas^6–10^. Based on this evidence, it has been proposed that memory is stored in a unified engram complex: a coordinated network of engram ensembles that spans anatomically distinct regions and functions collectively to store and retrieve specific memories^11–13^. In recent years, studies focusing on fear memory have enabled detailed characterization of memory processing, revealing the involvement of multiple brain regions, including the hippocampus^14–16^, amygdala^17,18^, and prefrontal cortex^19–21^, key centres of contextual fear memory function. In contrast, the neural correlates of episodic memory across the brain remain elusive. Episodic memory is defined as the recollection of past experiences bound to their spatial and temporal context^22,23^. The core features of episodic memory are conserved across mammalian and avian species, supported by anatomically and functionally homologous neural circuits^24^. The acquisition and recollection of episodic memories critically depend on the activity of the medial temporal lobe (MTL) system^25,26^, with the hippocampus playing a central role in binding spatial, temporal, and contextual information. Within this system, the entorhinal cortex (EC) serves as a central hub for the convergence of multimodal sensory and highly processed unimodal inputs, which are subsequently transmitted to the hippocampal formation to support the encoding of episodic components into unified memory representations^27^. Recently, growing evidence has highlighted the lateral sub-division of the entorhinal cortex (LEC) as a key region involved in processing contextual and temporal information. The LEC plays a central role in encoding associations between objects, places, and contexts, a core function in the formation of episodic-like memory^28–30^. Additionally, using a dual viral system based on targeted recombination in active populations (TRAP), it has been demonstrated that neuronal ensembles in the LEC support the encoding and retrieval of object-place-context associations, pointing to the existence of LEC engrams for these memories^31,32^. However, the specific contributions of brain regions outside the medial temporal lobe that participate in the encoding, storage, and retrieval of episodic memory remain poorly characterized. In recent years, advances in histological and imaging techniques, combined with pharmacological or behavioral induction of immediate early gene (IEG) expression, have enabled a comprehensive characterization of neuronal population activity across the whole brain. Brain-wide activity mapping approaches have been applied to a variety of paradigms, extending our knowledge of neural representations beyond individual brain regions^6,11,33,34^.

Starting from the hypothesis that episodic-like memory is encoded by distinct neuronal populations distributed across anatomically separate brain regions, we employed a brain-wide approach to map the activation of neuronal populations during recall of object-place-context recognition (OPC) memory, using the IEG c-Fos as a marker of recent neuronal activity. We then quantified c-Fos expression patterns across 75 brain regions, comparing episodic memory recall with three different control conditions and identified a subset of regions that were strongly activated during OPC memory recall. Since neuronal activation during memory recall alone is insufficient to establish the functional role of specific neuronal populations^3,35^, we next investigated the contribution of the identified neuronal populations to OPC memory recall. To this end, we used a TRAP-based viral strategy combined with chemogenetics^19,31^ to achieve tagging of learning-activated neurons and subsequent functional modulation of these neurons during memory retrieval. Finally, we recorded population activity during the recall of OPC memory from selected cortical regions, to investigate how these areas represent novel and familiar objects in freely behaving mice. Our findings suggest that episodic-like memory in mice depends on a fronto-posterior medial network, in which the retrosplenial cortex (RSP) emerges as a critical region for successful memory retrieval and contains neurons with properties of engram cells. The other regions identified within this network are necessary for memory expression but are unlikely to store the memory trace itself.

## Results

### Brain-wide c-Fos mapping reveals predominant isocortical activation during episodic-like memory recall

Based on the hypothesis that episodic-like memory is encoded and stored across multiple neuronal populations throughout the brain, we examined whether memory recall relies on a distributed network of interacting brain regions using the novel Object-Place-Context Recognition Task (OPCRT), a well-established modified version of the spontaneous recognition paradigm used to assess rodents’ ability to encode and retrieve object-place-context (OPC) associations^28,36^. To this end, we induced behavioral activation under different conditions and collected the brains 90 minutes later to capture the peak of c-Fos protein expression. To characterize the activation patterns associated with memory retrieval, while excluding areas involved in other aspects of exploration, we compared our experimental group subjected to the OPCRT with three control conditions: home-cage control (HC), context-exposed control (CNTX), and a group exposed to the same pair of objects across all contexts (OCT) (Figure **1A**). The collected brains were serially sectioned to obtain a coronal representation of the entire brain and subjected to immunohistochemical staining for c-Fos. Fluorescence images were then acquired using confocal microscopy and registered to the Allen Brain Atlas as previously described^37^. This process produced a comprehensive dataset of c-Fos^+^ cells across multiple levels of granularity, from major brain subdivisions to fine subregions, and across different behavioral conditions (Figure **1B**). Using PCA followed by k-means clustering, we assessed whether the c-Fos dataset could discriminate between the different behavioral groups. The analysis confirmed that the c-Fos activation profiles were sufficient to accurately distinguish all conditions. Inspection of the contributions of individual brain regions revealed that isocortical areas were the dominant contributors across the first three components (Figure **1C**). This suggests that isocortical regions critically shape the activation profiles that differentiate behavioral conditions. We then examined activation patterns across the eleven major brain subdivisions defined by the Allen Brain Atlas (Figure **1D**). As expected, groups subjected to behavioral activation exhibited stronger c-Fos expression compared to HC controls, reflecting the widespread recruitment of neurons during task performance. This effect was particularly pronounced in regions critically involved in sensory processing, memory, and integrative functions, including the isocortex, olfactory areas, hippocampal formation, cortical subplate, pallidum, and hypothalamus (Figure **1E** and **S1A**). To exclude the possibility that differences in c-Fos density were driven by variations in behavioral or locomotor parameters, we analyzed the performance of the three groups subjected to behavioral activation. As expected, the OPCRT group showed a significantly higher discrimination index and spent more time exploring the novel object, whereas the OCT group displayed no preference between the identical object pair (Figure **S1B**). Importantly, the total exploration time, number of interactions, and total distance travelled during the recall phase did not differ between the groups (Figure **S1C**, **S1D** and **S1E**).

**Figure 1:**
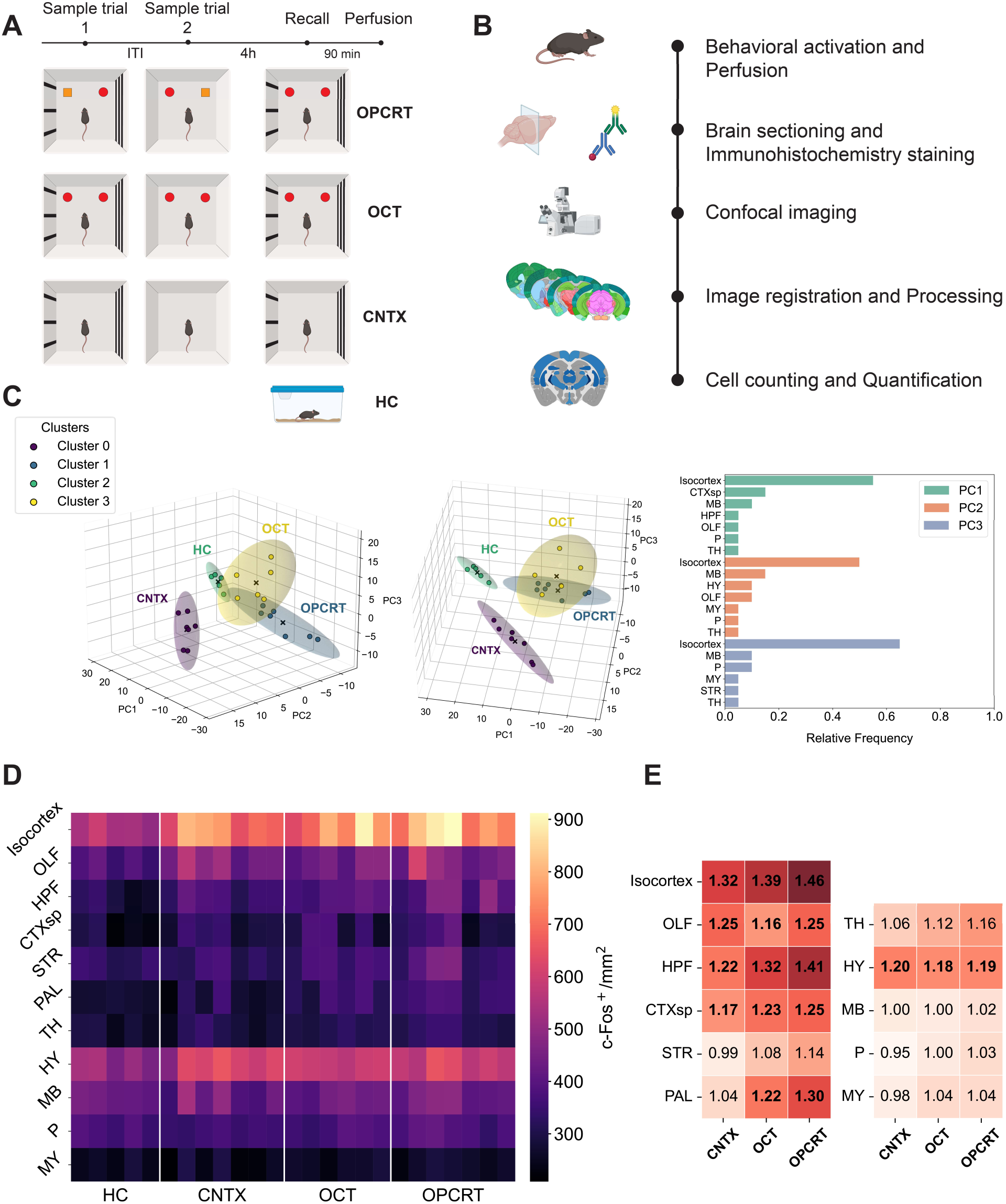
Brain-wide mapping of c-Fos activation during episodic-like memory recall. **(A)** Schematic representation of the behavioral paradigm used. Mice were assigned to four experimental groups: home cage (HC, n=5), context exposure (CNTX, n=7), object-context task (OCT, n=6), and object-place-context recognition task (OPCRT, n=7). In all conditions, both the sample trials and the recall phase lasted 10 min. **(B)** Overview of the experimental phases of the brain-wide c-Fos mapping. **(C)** Three-dimensional PCA representation of the c-Fos density dataset based on the first three principal components, with clusters identified by k-means clustering. Each dot represents an individual mouse. Two views of the same projection are shown from different angles. Right, analysis of the contribution of individual brain regions to each principal component showed a predominant involvement of isocortical areas across all three components. **(D)** Heatmap of c-Fos density across the eleven major brain subdivisions, displayed for individual mice across the four behavioral conditions. **(E)** Heatmap showing c-Fos fold change for each brain major subdivision in behaviorally activated groups (CNTX, OCT and OPCRT) relative to the HC condition. Fold-change values shown in bold are significantly different from HC, as assessed by two-way ANOVA followed by two-stage FDR post hoc correction (Q = 0.05; Isocortex: CNTX 1.323 ± 0.05 p < 0.001, OCT 1.389 ± 0.07 p < 0.001, OPCRT 1.458 ± 0.061 p < 0.001; OLF: CNTX 1.246 ± 0.058 p < 0.001, OCT 1.163 ± 0.047 p = 0.0311, OPCRT 1.253 ± 0.071 p < 0.001; HPF: CNTX 1.220 ± 0.047 p = 0.0028, OCT 1.321 ± 0.037 p < 0.001, OPCRT 1.406 ± 0.084 p < 0.001; CTXsp: CNTX 1.172 ± 0.052 p = 0.0189, OCT 1.230 ± 0.086 p = 0.0025, OPCRT 1.252 ± 0.080 p < 0.001; STR: CNTX 0.990 ± 0.034 p = 8929, OCT ± 1.078 ± 0.034 p = 0.0025, OPCRT 1.139 ± 0.073 p < 0.001; PAL: CNTX 1.038 ± 0.062 p = 0.5976, OCT 1.220 ± 0.051 p = 0.0038, OPCRT 1.296 ± 0.034 p < 0.001; TH: CNTX 1.059 ± 0.048 p = 0.4197, OCT 1.117 ± 0.021 p = 0.1217, OPCRT 1.163 ± 0.032 p = 0.0261; HY: CNTX 1.202 ± 0.046 p = 0.0058, OCT 1.184 ± 0.012 p = 0.0153, OPCRT 1.191 ± 0.028 p = 0.0092; MB: CNTX 0.995 ± 0.059 p = 0.9503, OCT 0.997 ± 0.027 p = 0.9918, OPCRT 1.022 ± 0.036 p = 0.6662; P: CNTX 0.951 ± 0.013 p = 4980, OCT 0.999 ± 0.028 p = 0.9918, OPCRT 1.031 ± 0.030 p = 0.6662; MY: CNTX 0.983 ± 0.054 p = 0.8150, OCT 1.036 ± 0.028 p = 0.6361, OPCRT 1.035 ± 0.029 p = 0.6288). Data are presented as mean ± SEM.

### Recruitment of a fronto-posterior medial network during episodic-like memory retrieval

To investigate potential variations in brain activation at a finer anatomical scale, we focused on a subset of 75 aggregated brain regions (Figure **2A**). To obtain a statistically robust comparison of activation patterns between the OPCRT group and control groups, we performed a mean-centered task partial least squares (PLS) analysis^38,39^. The comparison between HC and OPCRT revealed a widespread pattern of significantly different brain regions, indicating that the behavioral stimulation successfully induced a broad increase in c-Fos protein expression. Focusing on the isocortical regions, significant activation was observed in sensory regions such as the primary and secondary somatosensory cortices (SS) and the visual cortex (VIS). Frontal associative areas showing robust c-Fos expression comprised the anterior cingulate area (ACA), prelimbic (PL), infralimbic (ILA), and orbital (ORB) cortices, as well as the agranular insular cortex (AI). Additionally, higher-order associative regions such as the retrosplenial cortex (RSP), temporal association area (TEa), perirhinal cortex (PERI), and ectorhinal cortex (ECT) were strongly activated. Within the hippocampal formation and parahippocampal areas, significant c-Fos activation was observed in

**Figure 2:**
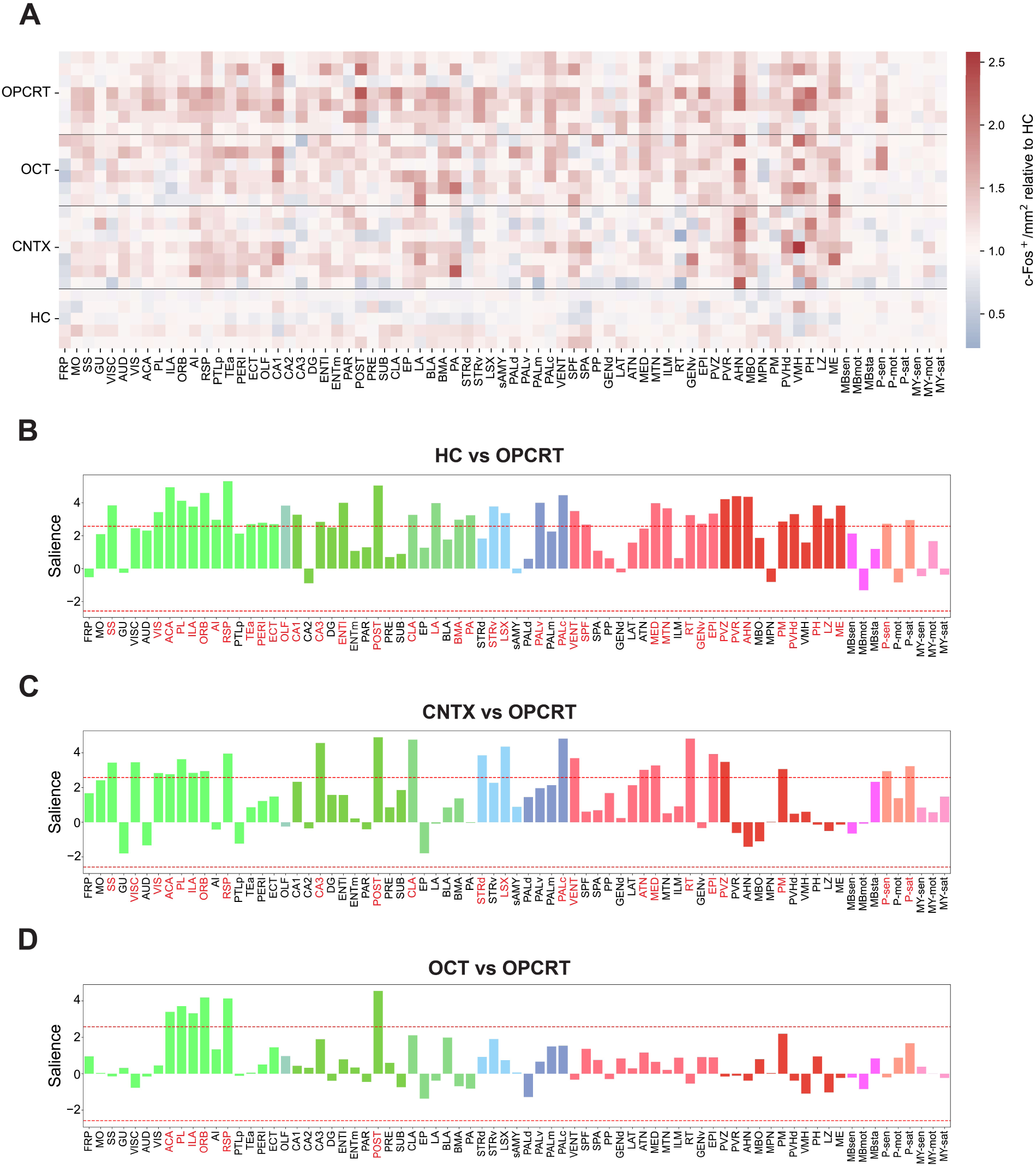
c-Fos activation at the aggregated mesoscale level. **(A)** Heatmap showing c-Fos density across 75 aggregated brain regions at the mesoscale level for individual mice across the four behavioral conditions. c-Fos density is shown relative to the HC condition. **(B-D)** Partial least squares (PLS) analysis of c-Fos expression comparing the OPCRT group with control conditions: HC **(B)**, CNTX **(C)**, and OCT **(D)**. Salience scores, normalized by the standard deviation estimated via bootstrap, identify brain regions that contribute most strongly to condition separation. Red dashed lines indicate the statistical threshold at salience scores of ± 2.58 (p < 0.01). Brain regions with salience scores exceeding the threshold are shown in red (Task PLS significant salience scores: HC vs OPCRT: VIS 3.418, ACA 4.801, PL 3.967, ILA 3.786, ORB 4.621, AI 2.970, RSP 5.329, TEa 2.696, PERI 2.716, ECT 2.689, OLF 3.866, CA1 3.331, CA3 2.851, ENTl 3.928, POST 5.064, CLA 3.258, LA 3.981, BMA 2.983, PA 3.188, STRv 3.796, LSX 3.449, PALv 4.008, PALc 4.479, VENT 3.559, SPF 2.659, MED 3.828, MTN 3.664, RT 3.254, GENv 2.719, EPI 3.363, PVZ 4.186, PVR 4.424, AHN 4.356, PM 2.845, PVHd 3.277, PH 3.760, LZ 3.008, ME 3.834, P-sen 2.678, P-sat 2.948; CNTX vs OPCRT: SS 3.453, VISC 3.464, VIS 2.866, ACA 2.790, PL 3.696, ILA 2.859, ORB 2.968, RSP 4.006, CA3 4.593, POST 5.003, CLA 4.768, STRd 3.884, LSX 4.367, PALc 4.790, VENT 3.760, ATN 3.047, MED 3.276, RT 4.876, EPI 3.999, PVZ 3.483, PM 3.040, P-sen 3.016, P-sat 3.246; OCT vs OPCRT: ACA 3.334, PL 3.684, ILA 3.328, ORB 4.218, RSP 4.144, POST 4.48).

CA1 and CA3 subfields, as well as in the lateral entorhinal cortex (ENTl). A strong activation was observed in the postsubiculum (POST) and claustrum (CLA), key regions involved in spatial processing and cortical integration. The remaining significantly activated regions primarily reflect engagement of olfactory, limbic, thalamic, and hypothalamic circuits, suggesting a broad network of subcortical and sensory structures involved in spatial navigation, object interaction, and memory recall (Figure **2B**). By comparing OPCRT to CNTX groups, we were able to disentangle memory-specific activation from the baseline activity related to general context exploration (Figure **2C**). This analysis revealed that primary sensory and frontal associative cortical areas exhibited a significantly higher density of c-Fos^+^ neurons, supporting their involvement in object exploration and memory retrieval. Within the hippocampal formation, significant activation was detected in CA3 and the POST. In addition, we observed recruitment of the claustrum, along with more selective activation of subcortical structures previously identified in the comparison with the HC control. Finally, when comparing the OPCRT group with the control condition sharing the most similar sensory input (OCT), a strong reduction in the number of significantly activated regions was observed. The pattern of c-Fos activation between the two groups was largely similar, with notable exceptions in subregions of the medial prefrontal cortex (ACA, PL, ILA), the ORB, the RSP, and the POST; resulting in a subset of regions critically involved in executive function and decision-making (mPFC)^40,41^, value representation and flexible behavior (ORB)^42^, spatial navigation and contextual processing (RSP)^43,44^, spatial orientation and memory integration (POST)^45^ (Figure **2D**). These comparisons indicate a progressive restriction of recruited brain regions associated with increasing cognitive complexity. In fact, the spatial distribution of c-Fos fold change across coronal brain sections (Figure **3A**), which reveals widespread activation relative to HC that becomes progressively confined when comparisons are restricted to CNTX and OCT controls, as confirmed by the nested structure of the Venn diagram showing significantly activated brain regions across behavioral conditions (Figure **3B**). The core component of this structure, comprising the mPFC, RSP, ORB, and POST, identifies brain regions that are robustly and consistently engaged during episodic-like memory recall (Figure **3C**). To explore the anatomical organization of the activated brain regions, we constructed an anatomical connectivity network using data from the Allen Brain Connectivity Atlas^46^. We focused on direct connections among the activated regions, based on the hypothesis that co-activated areas may form a dedicated subnetwork with preferential anatomical connectivity. We observed two highly connected divisions composed of frontal and posterior medial areas (Figure **3D**). Interestingly, this subnetwork exhibited significantly higher mean connectivity compared to equivalent random networks, suggesting that it constitutes a specialized anatomical unit supporting novelty-driven behavior (Figure **3E**).

**Figure 3:**
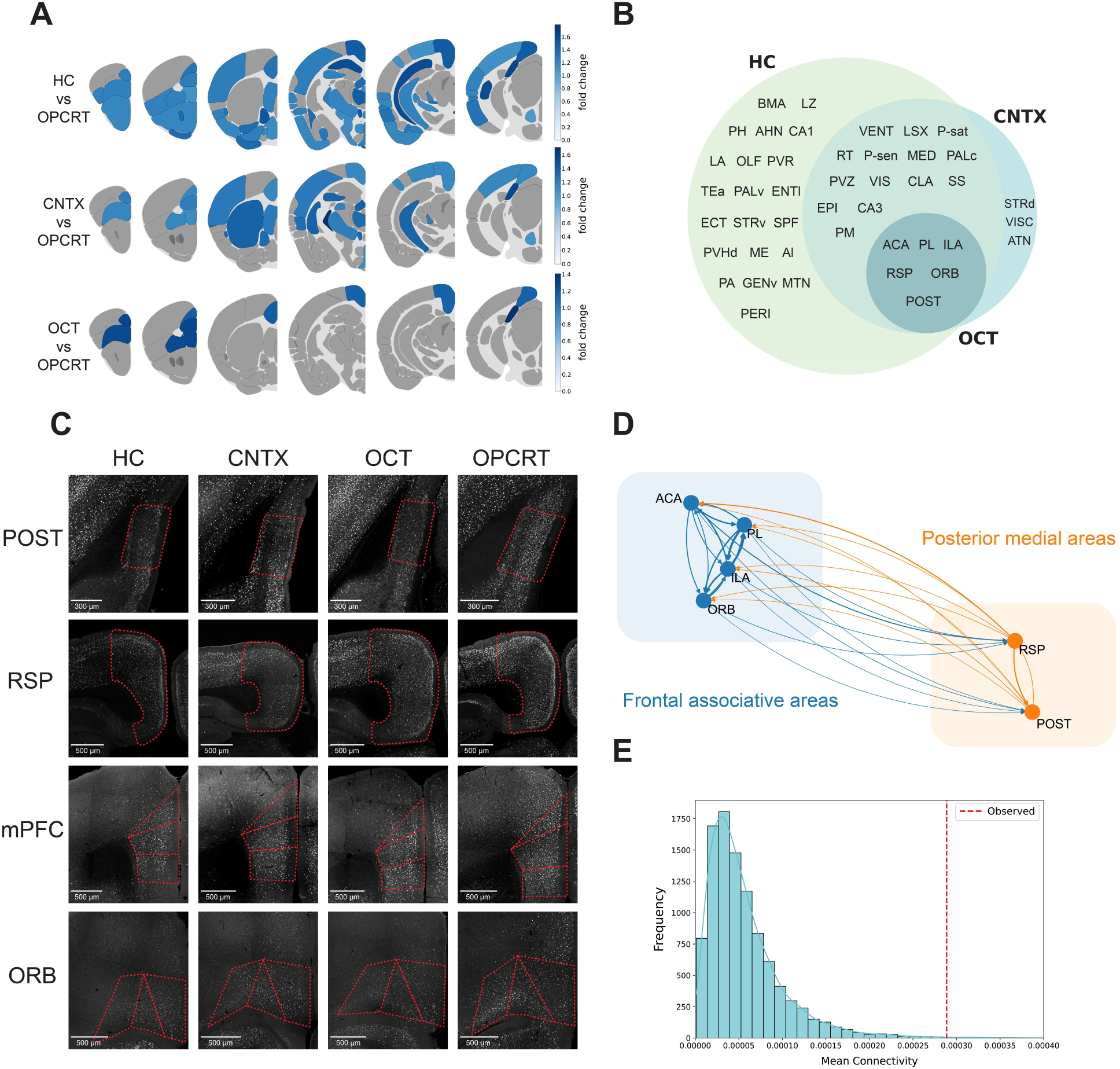
Brain-wide c-Fos mapping reveals a core episodic-like memory network. **(A)** Brain-wide heatmaps showing c-Fos density fold-change during OPC memory recall relative to each control condition (HC, CNTX, and OCT). Only significantly activated brain regions are colored and mapped across coronal brain sections at the aggregated mesoscale level. **(B)** Venn diagram summarizing brain regions showing significantly higher activation in the OPCRT group compared to each of the other conditions, as identified by task PLS. **(C)** Representative c-Fos immunohistochemistry for the four brain regions significantly activated relative to the OCT group (mPFC, RSP, ORB, and POST), shown across all experimental groups. Brain region boundaries are indicated by red dashed outlines; scale bars are shown in the images. **(D)** Anatomical network composed of areas significantly activated relative to the OCT group, with brain regions shown as nodes and directed anatomical connections represented as arrows. The network reveals two major divisions corresponding to frontal associative and posterior medial components. **(E)** Distribution of mean connectivity of equivalent random anatomical networks obtained by bootstrapping (n = 10000). Mean connectivity represents the average normalized projection density between region pairs. The red dashed line indicates the mean connectivity of the subnetwork composed of brain regions activated relative to the OCT group. The observed value was significantly greater than values obtained from equivalent random networks (observed mean = 0.00029; random mean = 0.00006; one-sided permutation test, **p = 0.0027).

### Network analysis indicates a shift toward a more efficient network topology during memory recall

While the quantification of c-Fos expression provides a map of activated brain areas, it does not capture how these regions operate as an integrated system. To address this limitation, we employed functional connectivity analysis to assess how regional activity co-varies across animals following behavioral activation. To estimate patterns of functional connectivity, we computed interregional correlation matrices for each experimental group subjected to behavioral activation (CNTX, OCT, and OPCRT) at the aggregated level of anatomical granularity. Pearson correlation matrices were then thresholded by retaining only the strongest statistically significant correlations (r > 0.80; p < 0.05), from which we generated functional networks in which nodes represent individual brain regions and edges represent suprathreshold functional connections (Figure **4A**, **S2A** and **S2B**). To further characterize the topological organization of the resulting functional networks, we quantified a set of well-established graph metrics^47^. Compared to CNTX and OCT networks, the OPCRT network exhibited a significant increase in node degree and overall network density (Figure **4B** and **4C**), indicating a more densely interconnected structure. In parallel, global efficiency was significantly elevated in the OPCRT network (Figure **4D**), suggesting that information transfer across the network can occur through shorter and more efficient paths. At the local scale, the OPCRT network showed an increased clustering coefficient (Figure **4E**), consistent with tighter local connectivity and a more pronounced modular organization. In contrast, betweenness centrality did not increase in the OPCRT network relative to the OCT network, indicating no shift toward greater network centralization (Figure **4F**). Finally, the OPCRT network exhibited a higher small-world index (Figure **4G**), a topological profile that optimally balances local specialization with global integration. Taken together, these alterations in topological parameters indicate that memory retrieval in the OPCRT condition was associated with a more densely connected and efficient network organization.

**Figure 4.**
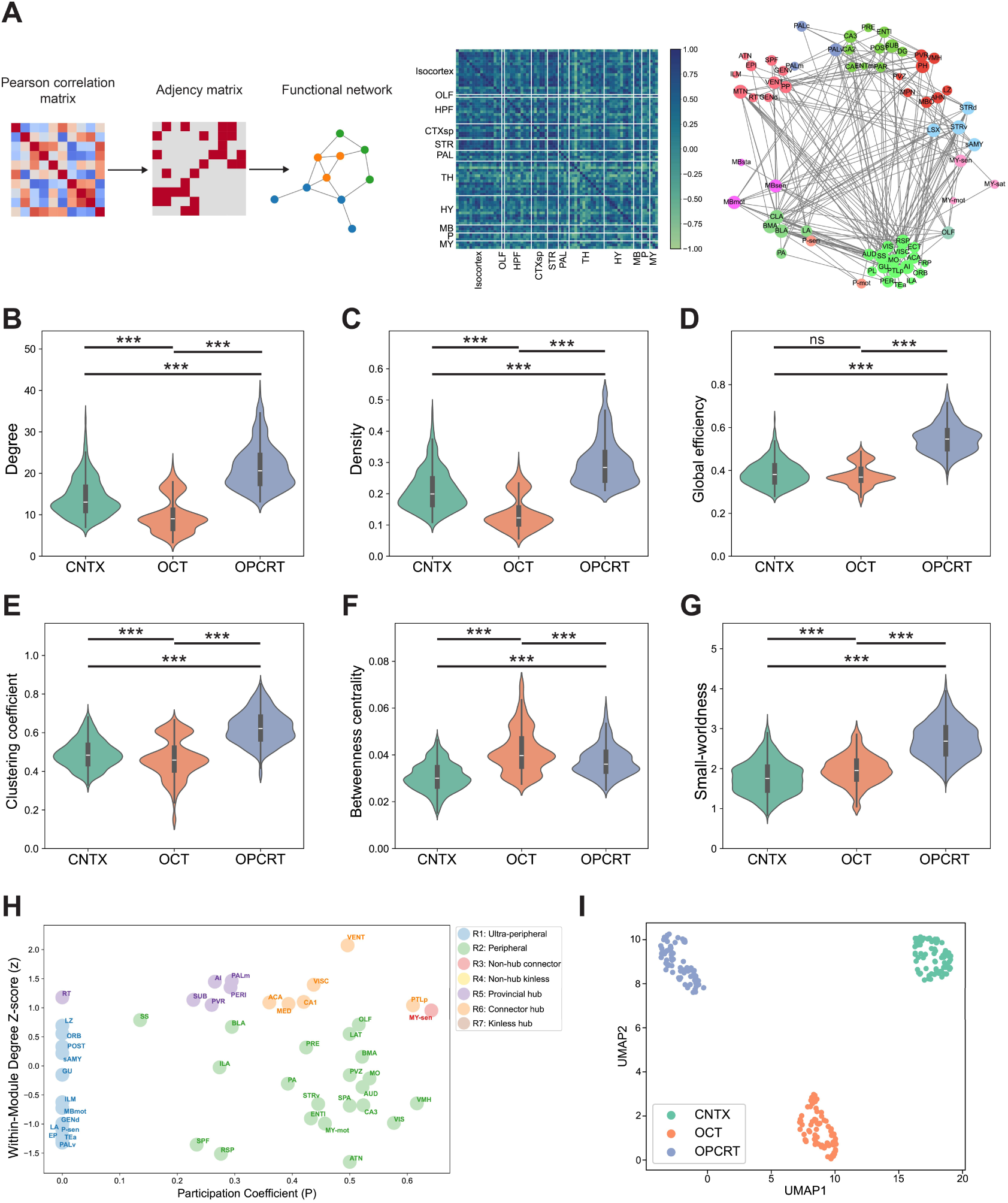
Topological analysis of functional brain networks: **(A)** Representative image of the functional connectivity analysis for the OPCRT group. For each condition, the unfiltered interregional Pearson correlation matrix computed across aggregated mesoscale brain regions is shown on the left. Major anatomical subdivisions are indicated along the matrix axes. On the right, the corresponding functional network is shown, obtained by retaining only positive correlations exceeding r > 0.8 that were statistically significant (p < 0.05). Nodes represent individual brain regions and are colored by major anatomical subdivision, while edges represent suprathreshold functional connections. Topological properties of functional networks were estimated using bootstrap resampling across subjects. Functional networks were recomputed over 1000 bootstrap iterations, and graph metrics were calculated for each iteration to obtain distributions of network topology measures. **(B)** Node degree was significantly higher in the OPCRT group compared to CNTX and OCT (Kruskal-Wallis test with Dunn’s multiple comparisons test: CNTX 14.16 ± 0.143 vs OCT 9.912 ± 0.136, *** p < 0.001; CNTX vs OPCRT 21.50 ± 0.167, *** p < 0.001; OCT vs OPCRT, *** p < 0.001). **(C)** Network density was significantly increased in OPCRT relative to both control conditions (Kruskal-Wallis test with Dunn’s multiple comparisons test: CNTX 0.2127 ± 0.002 vs OCT 0.138 ± 0.001, *** p < 0.001; CNTX vs OPCRT 0.299 ± 0.002, *** p < 0.001; OCT vs OPCRT, *** p < 0.001). **(D)** Global efficiency was significantly higher in OPCRT compared to CNTX and OCT (Kruskal-Wallis test with Dunn’s multiple comparisons test: CNTX 0.388 ± 0.001 vs OCT 0.379 ± 0.001, ns p = 0.0859; CNTX vs OPCRT 0.550 ± 0.002, *** p < 0.001; OCT vs OPCRT, *** p < 0.001). **(E)** Clustering coefficient was significantly increased in OPCRT relative to both control groups (Kruskal-Wallis test with Dunn’s multiple comparisons test: CNTX 0.490 ± 0.002 vs OCT 0.464 ± 0.003, *** p < 0.001; CNTX vs OPCRT 0.627 ± 0.002, *** p < 0.001; OCT vs OPCRT, *** p < 0.001). **(F)** Betweenness centrality in OPCRT was significantly higher than CNTX but lower than OCT (Kruskal-Wallis test with Dunn’s multiple comparisons test: CNTX 0.030 ± 0.0001 vs OCT 0.041 ± 0.0003, *** p < 0.001; CNTX vs OPCRT 0.037 ± 0.0002, *** p < 0.001; OCT vs OPCRT, *** p < 0.001). **(G)** Small-worldness was significantly higher in OPCRT compared to CNTX and OCT (Kruskal-Wallis test with Dunn’s multiple comparisons test: CNTX 1.777 ± 0.013 vs OCT 1.975 ± 0.011, *** p < 0.001; CNTX vs OPCRT 2.714 ± 0.014, *** p < 0.001; OCT vs OPCRT, *** p < 0.001). **(H)** Identification of hub regions and topological roles within the OPCRT functional network. Nodes were classified based on their within-module degree z-score and participation coefficient, allowing assignment to specific topological roles. **(I)** UMAP embedding of interregional functional connectivity profiles. Each point represents a brain region within a given condition, positioned according to its correlation pattern. Brain regions are colored by behavioral conditions, revealing a clear segregation of CNTX, OCT, and OPCRT networks based on distinct patterns of c-Fos co-activation. Data are presented as mean ± SEM.

To further investigate the internal organization of the functional networks identified across conditions, we next focused on the identification of hub regions (Figure **4H**, **S2C** and **S2D**). To characterize the contribution of each brain region to network organization, nodes were classified based on their within-module connectivity and their participation coefficient^48^. Focusing on the OPCRT network, the POST, RSP, SUB, and posterior hypothalamic nucleus (PH) exhibited connector hub (R6) characteristics, indicating a central role in mediating communication between distinct network modules. In contrast, several regions, including perirhinal cortex (PERI), auditory cortex (AUD), anterior hypothalamic nucleus (AHN), lateral entorhinal cortex (ENTl), lateral amygdala (LA), ectorhinal cortex (ECT), posterior parietal association area (PTLp), and somatosensory cortex (SS), were classified as provincial hubs (R5). These regions displayed high within-module connectivity, consistent with a role in supporting locally specialized processing in individual functional modules rather than inter-module integration. It is also worth noting that the VIS and DG displayed non-hub connector (R3) properties. Although these regions did not reach hub-level connectivity, their elevated participation coefficients indicate that they contribute to linking multiple network modules. The attribution of these topological roles, together with the observation that RSP and POST are significantly activated during episodic-like memory retrieval, emphasizes the contribution of the posterior medial component of the proposed anatomical network, suggesting that the functional network is organized around a set of posterior medial integrative nodes that coordinate interactions between different modules during episodic-like memory recall. To further assess whether each behavioral condition is associated with a distinct pattern of interregional co-activation, we applied a low-dimensional embedding approach using Uniform Manifold Approximation and Projection (UMAP). In this embedding, each brain region was represented by its pattern of correlations with all other regions, and these high-dimensional profiles were projected into a two-dimensional space such that regions with similar correlation patterns were positioned closer together. The resulting UMAP representation revealed a clear segregation of regions according to behavioral conditions, indicating that each condition is characterized by a distinct configuration of functional interactions of c-Fos activity (Figure **4I**).

### Dissecting the role of learning-tagged neurons during episodic memory retrieval using chemogenetics

Although significant activation of specific neuronal populations suggests their engagement in the network supporting memory retrieval, such evidence alone is insufficient to define these populations as memory engrams. To causally assess the functional contribution of brain regions identified as activated during OPCRT recall, we employed a TRAP-based dual-virus strategy to selectively express chemogenetic receptors during memory encoding^19^. The activity of the learning-tagged neuronal population was subsequently manipulated using CNO during memory recall, either at 12h to induce inhibition, when memory expression is still intact, or at 48h to induce activation, when the memory is no longer behaviorally expressed under natural recall conditions (Figure **5A**). Using this approach, we examined the contribution of selected brain regions to memory recall, focusing on the POST, RSP, mPFC and ORB (Figure **5B** and **5C**). We first targeted the mPFC (ACA, PL, and ILA) by expressing chemogenetic receptors in learning-tagged neurons using the TRAP system, enabling either inhibition or activation of this subpopulation. Control mice tested at 12h exhibited the expected preference for the object in the novel configuration, whereas control mice tested at 48h showed no preference, indicating that OPC memory is no longer accessible at this time point. Chemogenetic inhibition of learning-tagged mPFC neurons resulted in a significant impairment of memory performance when mice were tested at 12h, suggesting that the activity of this neuronal population is necessary for successful memory retrieval. In contrast, chemogenetic reactivation of the same population at 48h did not facilitate memory recall, indicating that activation of mPFC learning-tagged neurons is not sufficient to restore memory expression once natural retrieval has decayed (Figure **5D** and **5E**). The same chemogenetic approach applied to the RSP resulted in a similar significant impairment of memory performance at 12h. In addition, chemogenetic reactivation of this neuronal population at 48h significantly facilitated memory recall, a time point at which natural cues alone were insufficient to support memory expression (Figure **5F** and **5G**). Similarly to the mPFC, chemogenetic manipulation of learning-tagged neurons in the ORB (Figure **5H** and **5I**) and POST (Figure **5J** and **5K**) impaired memory performance at 12h but failed to facilitate memory recall at 48h. Importantly, for all the areas investigated, changes in memory performance were not accompanied by alterations in locomotor activity or in the number of object interactions (Figure **S3A**-**S3H**).

**Figure 5:**
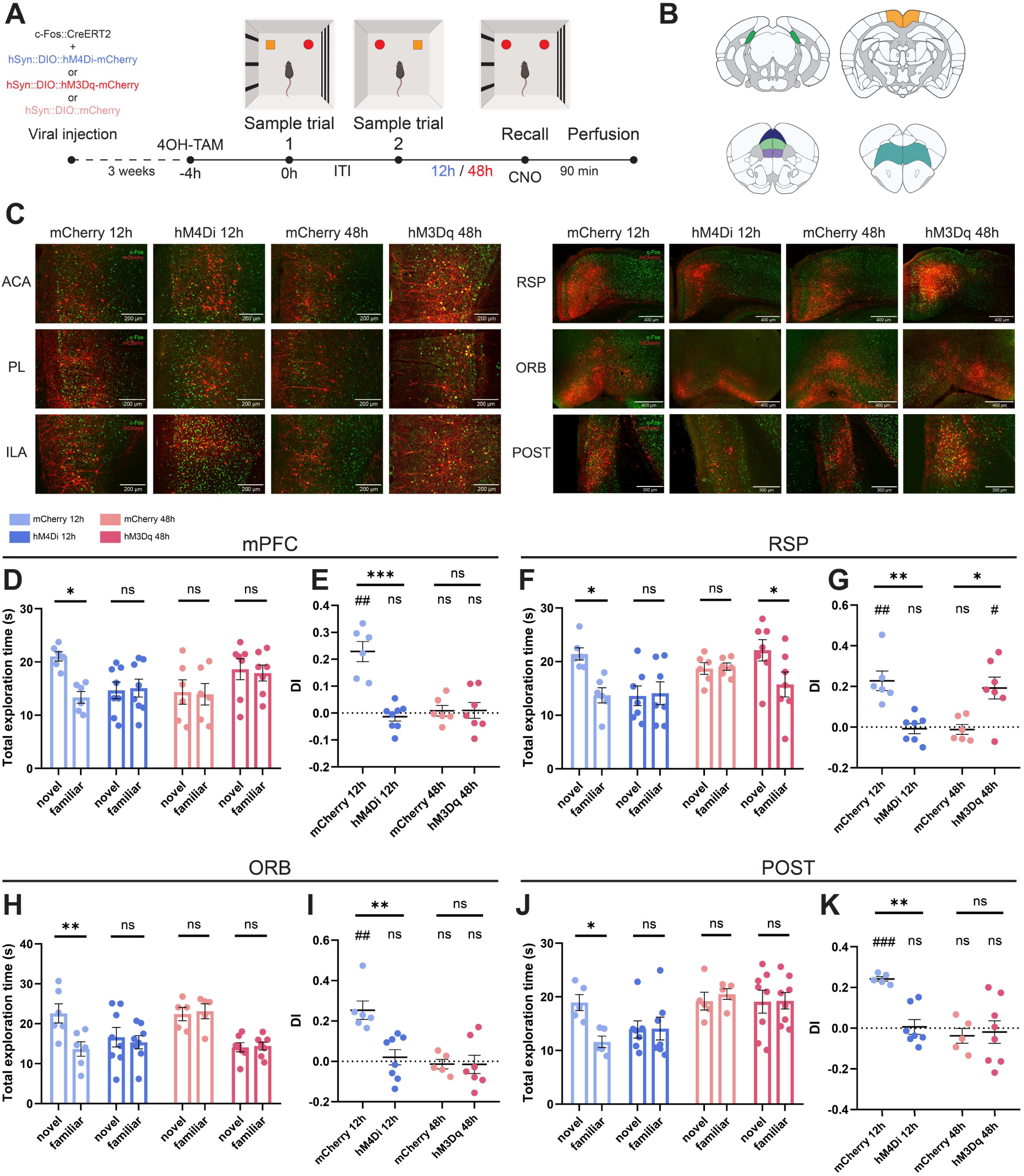
Chemogenetic manipulation of learning-tagged neuronal populations during OPC memory retrieval. **(A)** Schematic representation of the TRAP-based chemogenetic manipulation during the OPCRT. The AAV vectors used for neuronal tagging and chemogenetic manipulation are shown on the left. Neuronal populations activated during OPC learning were tagged by 4OH-TAM administration and subsequently manipulated during the recall phase using CNO, either at 12h to induce inhibition or at 48h to induce activation. Mice were perfused 90 min after the recall session. **(B)** Coronal brain schematics illustrating the anatomical location of the regions targeted for chemogenetic manipulation, including the postsubiculum (POST), retrosplenial cortex (RSP), medial prefrontal cortex (mPFC), and orbitofrontal cortex (ORB). **(C)** Representative c-Fos immunohistochemistry of the targeted brain regions. The mPFC is shown separately for its three subregions: anterior cingulate cortex (ACA), prelimbic cortex (PL), and infralimbic cortex (ILA). Images are shown for the mCherry 12h, hM4Di 12h, mCherry 48h, and hM3Dq 48h groups. mCherry-expressing neurons are shown in red and c-Fos^+^ cells in green. Scale bars are indicated in the images. For the mPFC, analysis of the total object exploration time **(D)** during the recall phase revealed differential preference for the novel association across experimental groups. A significant increase in exploration of the novel relative to the familiar configuration was observed in the mCherry 12h group, whereas this preference was impaired following chemogenetic inhibition at 12h. No significant difference in exploration time between novel and familiar configuration was detected in either group tested at 48h (two-way ANOVA with Sidak’s multiple comparisons test: mCherry 12h Novel 21.04 ± 0.85 n = 6 vs mCherry 12h Familiar 13.34 ± 1.08 n = 6, * p = 0.162; hM4Di 12h Novel 14.66 ± 1.57 n = 8 vs hM4Di 12h Familiar 15.08 ± 1.68 n = 8, ns p = 0.995; mCherry 48h Novel 14.34 ± 2.29 n = 6 vs mCherry 48h Familiar 13.92 ± 2.02 n = 6, ns p = 0.9997; hM3Dq 48h Novel 18.62 ± 1.97 n = 7 vs hM3Dq 48h 12h Familiar 17.91 ± 1.50 n = 7, ns p = 0.9969). **(E)** Discrimination index (DI) during the recall phase across experimental groups. DI values were significantly greater than chance level only in the mCherry 12h group (mCherry 12h 0.228 ± 0.037, n = 6, ** p = 0.0017, df = 5, t = 6.105; hM4Di 12h -0.013 ± 0.016, n = 8, ns p = 0.4418, df = 7, t = 0.8152; mCherry 48h 0.008 ± 0.019, n = 6, ns p = 0.6821, df = 5, t = 0.4344; hM3Dq 48h 0.009 ± 0.029, n = 7, ns p = 0.7529, df = 6, t = 0.3295; one-sample t test). A significant difference in DI was observed between the mCherry 12h and hM4Di 12h groups, whereas no significant difference was detected between the two groups tested at 48h (one-way ANOVA with Sidak’s multiple comparisons test: mCherry 12h vs hM4Di 12h, *** p < 0.001; mCherry 48h vs hM3Dq 48h, ns p = 0.9999). For the RSP, **(F)** mCherry 12h group showed a significant preference for the object in the novel configuration relative to the familiar one. Chemogenetic inhibition at 12h (hM4Di 12h) impaired novel object exploration, with no difference between novel and familiar exploration times. At 48h, no preference was observed in the mCherry group, whereas chemogenetic activation (hM3Dq 48h) resulted in significantly greater exploration of the novel object compared to the familiar one (two-way ANOVA with Sidak’s multiple comparisons test: mCherry 12h Novel 21.44 ± 1.15 n = 6 vs mCherry 12h Familiar 13.72 ± 1.41 n = 6, * p = 0.0176; hM4Di 12h Novel 13.62 ± 1.83 n = 8 vs hM4Di 12h Familiar 14.11 ± 2.13 n = 7, ns p = 0.9993; mCherry 48h Novel 18.72 ± 1.07 n = 6 vs mCherry 48h Familiar 19.05 ± 0.67 n =6, ns p = 0.9999; hM3Dq 48h Novel 22.12 ± 1.98 n = 7 vs hM3Dq 48h 12h Familiar 15.73 ± 2.31 n = 7, * p = 0.0403). **(G)** DI values were significantly greater than 0 in the mCherry 12h and hM3Dq 48h groups, indicating a preference for the object in the novel configuration. In contrast, DI following chemogenetic inhibition at 12h did not differ from chance, indicating a loss of preference for the novel association (mCherry 12h 0.227 ± 0.048, n = 6, ** p = 0.0054, df = 5, t = 4.684; hM4Di 12h -0.007 ± 0.024, n = 7, ns p = 0.7687, df = 6, t = 0.3078; mCherry 48h -0.011 ± 0.024, n = 6, ns p = 0.6571, df = 5, t = 0.4716; hM3Dq 48h 0.1922 ± 0.053, n = 7, * p = 0.0118, df = 6, t = 3.572; one-sample t test). DI also differed significantly between each chemogenetic group and its respective control (one-way ANOVA with Sidak’s multiple comparisons test: mCherry 12h vs hM4Di 12h, ** p = 0.003; mCherry 48h vs hM3Dq 48h, * p = 0.0111). For the ORB, **(H)** the mCherry 12h group showed a significant preference for the object in the novel configuration. Chemogenetic inhibition at 12h (hM4Di 12h) resulted in a loss of preference for the novel association. At 48h, neither the mCherry nor the hM3Dq group showed a significant preference (two-way ANOVA with Sidak’s multiple comparisons test: mCherry 12h Novel 22.59 ± 2.40 n = 6 vs mCherry 12h Familiar 13.63 ± 1.81 n = 6, ** p = 0.008; hM4Di 12h Novel 16.62 ± 2.45 n = 8 vs hM4Di 12h Familiar 15.36 ± 1.63 n = 8, ns p = 0.9738; mCherry 48h Novel 22.41 ± 1.66 n = 5 vs mCherry 48h Familiar 23.11 ± 1.86 n =5, ns p = 0.9988; hM3Dq 48h Novel 14.09 ± 1.16 n = 7 vs hM3Dq 48h 12h Familiar 14.40 ± 0.99 n = 7, ns p = 0.9999). **(I)** The mCherry 12h group showed a DI significantly greater than chance, indicating a preference for the novel association. Chemogenetic inhibition at 12h reduced DI to chance level, and chemogenetic activation at 48h failed to restore memory performance group (mCherry 12h 0.253 ± 0.045, n = 6, ** p = 0.0026, df = 5, t = 5.565; hM4Di 12h -0.020 ± 0.037, n = 8, ns p = 0.6061, df = 7, t = 0.5397; mCherry 48h -0.013 ± 0.023, n = 5, ns p = 0.5947, df = 4, t = 0.5772; hM3Dq 48h -0.014 ± 0.045, n = 7, ns p = 0.7542, df = 6, t = 0.3279; one-sample t test). DI differed significantly between the mCherry 12h and hM4Di 12h groups, but not between the two groups tested at 48h (one-way ANOVA with Sidak’s multiple comparisons test: mCherry 12h vs hM4Di 12h, ** p = 0.002; mCherry 48h vs hM3Dq 48h, ns p = 0.9999). For the POST, **(J)** The mCherry 12h group showed a significant preference for the object in the novel configuration. Chemogenetic inhibition at 12h (hM4Di 12h) impaired novel object exploration. At 48h, neither the mCherry nor the hM3Dq group showed a significant preference (two-way ANOVA with Sidak’s multiple comparisons test: mCherry 12h Novel 18.92 ± 1.48 n = 5 vs mCherry 12h Familiar 11.60 ± 1.07 n = 5, * p = 0.0498; hM4Di 12h Novel 13.92 ± 1.61 n = 7 vs hM4Di 12h Familiar 14.08 ± 2.12 n = 7, ns p = 0.9999; mCherry 48h Novel 19.09 ± 2.13 n = 5 vs mCherry 48h Familiar 20.52 ± 0.99 n =5, ns p = 0.9831; hM3Dq 48h Novel 19.09 ± 2.13 n = 8 vs hM3Dq 48h 12h Familiar 19.27 ± 1.54 n = 8, ns p = 0.9999). **(K)** DI values were significantly greater than 0 only in the mCherry 12h group. Chemogenetic inhibition at 12h reduced DI to chance level, and chemogenetic activation at 48h did not facilitate memory recall (mCherry 12h 0.241 ± 0.011, n = 5, *** p < 0.001, df = 4, t = 21.81; hM4Di 12h 0.006 ± 0.036, n = 7, ns p = 0.8695, df = 6, t = 0.1715; mCherry 48h -0.037 ± 0.036, n = 5, ns p = 0.3647, df = 4, t = 1.022; hM3Dq 48h -0.019 ± 0.055, n = 8, ns p = 0.7392, df = 7, t = 0.3465; one-sample t test). DI differed significantly between the mCherry 12h and hM4Di 12h groups, but not between the two groups tested at 48h (one-way ANOVA with Sidak’s multiple comparisons test: mCherry 12h vs hM4Di 12h, ** p = 0.009; mCherry 48h vs hM3Dq 48h, ns p = 0.9999). Data are presented as mean ± SEM.

At the cellular level, the density of c-Fos^+^ cells across the selected brain regions did not differ following either inhibitory or excitatory chemogenetic manipulations. During natural memory recall, however, c-Fos density was significantly higher at 12h compared to 48h, confirming that recent memory recall was associated with elevated c-Fos activation in this specific circuit (Figure **6A**-**6D**). Analysis of ensemble reactivation revealed that chemogenetic inhibition reduced the proportion of reactivated cells, whereas chemogenetic stimulation induced c-Fos expression in nearly all tagged neurons. Notably, successful natural memory recall at 12h was associated with significantly higher ensemble reactivation in the mPFC, RSP, and POST, but not in the ORB, compared to 48h (Figure **6E**-**6H**). Consistent with this, ensemble reactivation in these same regions exceeded chance levels in the mCherry 12h group, but not in the mCherry 48h group. In the ORB, recent and late recall induced a similar proportion of reactivation, suggesting that activity in this region may be required for correct memory recall, but is not directly associated with memory performance. Interestingly, although at 48h the reactivation proportion in the mPFC and RSP did not differ significantly from chance levels, the POST still showed a trend toward significant ensemble activation (Figure **6I**-**6L**). No significant correlation was observed between behavioral performance and either the percentage of ensemble reactivation or c-Fos density (Figure **S3I**-**S3L**) in the selected brain regions, with the exception of the RSP that showed a positive correlation during successful memory recall at 12h (Figure **6M**-**6P**). Overall, these findings indicate that the RSP is strongly involved in successful episodic memory recall and contains a subpopulation of learning-tagged neurons with properties of engram cells. To further assess whether these effects differed across mPFC subregions, we next examined c-Fos expression, ensemble reactivation, and their relationship with behavioral performance separately within the ACA, PL, and ILA. Focusing on the ACA, we observed that c-Fos density was significantly higher in the 12h control group compared to the 48h control group. In contrast, the percentage of ensemble reactivation did not differ between these conditions, suggesting that both successful and unsuccessful memory recall engaged a similar proportion of the learning-tagged neurons in this subregion. No significant deviation from chance-level reactivation was observed in either group, indicating that ACA learning-tagged neurons may be recruited to a comparable extent regardless of memory performance. In fact, we did not observe a significant relationship between DI and either c-Fos expression or ensemble reactivation in this subregion (Figure **S4A**). In contrast to the ACA, both the PL and ILA exhibited a pattern closely resembling that observed across the entire mPFC. In both subregions, c-Fos^+^ density and the percentage of ensemble reactivation were significantly higher in the 12h control group compared to the 48h group. Ensemble reactivation exceeded chance levels only in the 12h group, while 48h group levels were not different from the chance. Interestingly, within the PL, we observed a strong positive correlation between memory performance and ensemble reactivation in the mCherry 12h group (Figure **S4B** and **S4C**). These findings suggest that distinct mPFC subregions differentially contribute to OPC memory retrieval, with PL and ILA ensembles showing reactivation patterns closely associated with memory accessibility, whereas ACA involvement appears to reflect a more general engagement during recall that is not accompanied by selective reactivation of learning-tagged ensembles.

**Figure 6:**
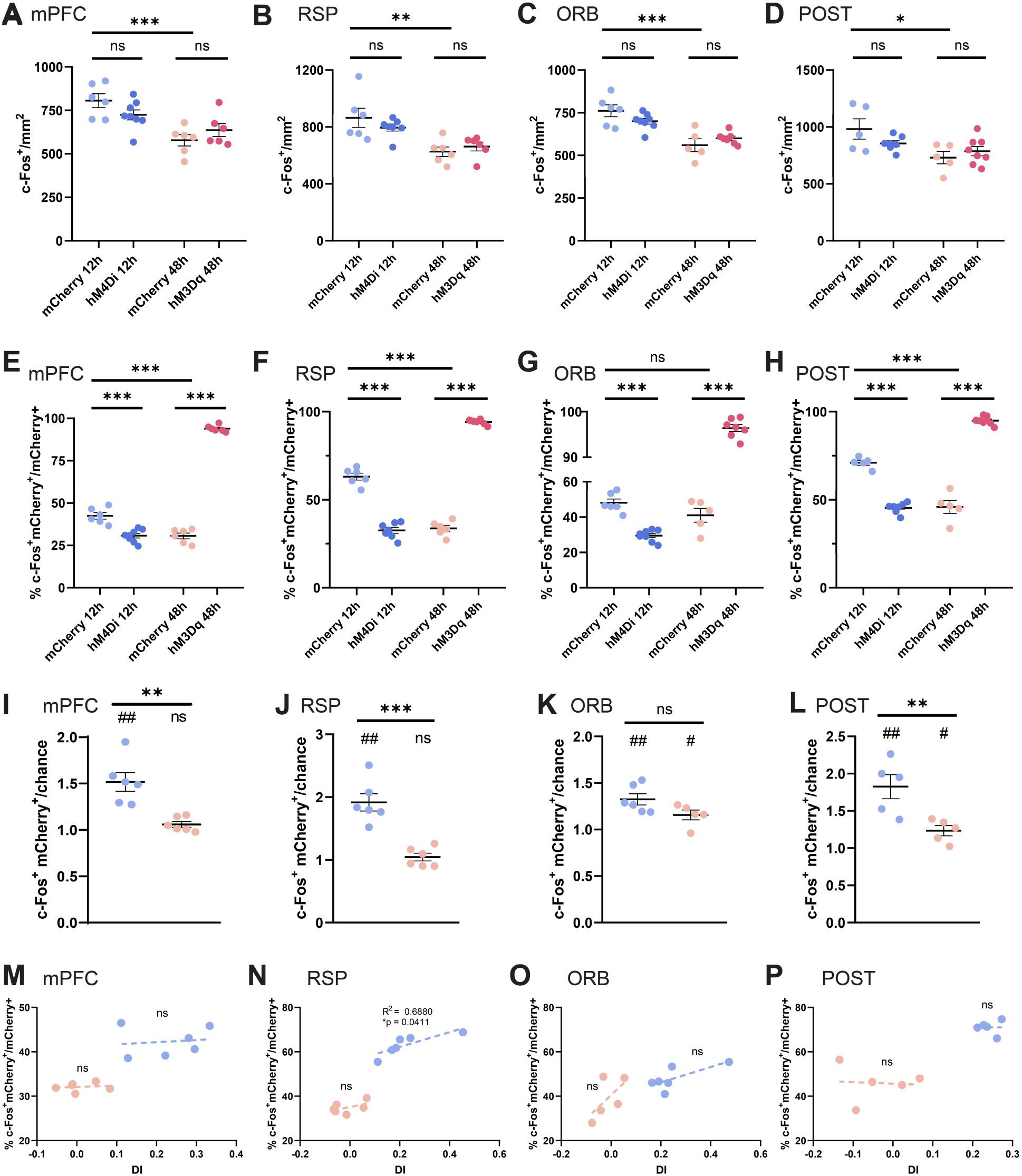
c-Fos density and ensemble reactivation across selected brain regions during OPC memory retrieval. Quantification of c-Fos^+^ cell density across the selected brain regions during OPC memory recall. **(A)** c-Fos density across the entire mPFC did not differ following inhibitory or excitatory chemogenetic manipulation. Under control conditions, c-Fos density was significantly higher at 12h compared to 48h (one-way ANOVA with Sidak’s multiple comparisons test: mCherry 12h 806.6 ± 39.26 n = 6 vs hM4Di 12h 724.4 ± 28.84 n = 8, ns p = 0.3204; mCherry 48h 578 ± 32.34 n = 6 vs hM3Dq 48h 636.4 ± 37.05 n = 6, ns p = 0.6530; mCherry 12h vs mCherry 48h, *** p < 0.001). **(B)** Quantification of c-Fos density in the RSP. c-Fos density did not differ between groups subjected to chemogenetic manipulation and their respective controls, but was significantly higher in the mCherry 12h group compared to the mCherry 48h group (one-way ANOVA with Sidak’s multiple comparisons test: mCherry 12h 864.2 ± 66.93 n = 6 vs hM4Di 12h 796 ± 24.85 n = 7, ns p = 0.6375; mCherry 48h 626.3 ± 33.57 n = 6 vs hM3Dq 48h 662.3 ± 30.44 n = 6, ns p = 0.9289; mCherry 12h vs mCherry 48h, ** p = 0.0033). In the ORB, **(C)** c-Fos density did not differ between chemogenetically manipulated groups and their respective controls, but was significantly higher in the mCherry 12h group compared to the mCherry 48h group (one-way ANOVA with Sidak’s multiple comparisons test: mCherry 12h 761.4 ± 34.71 n = 6 vs hM4Di 12h 700.4 ± 16.64 n = 8, ns p = 0.3061; mCherry 48h 560 ± 38.18 n = 5 vs hM3Dq 48h 600.2 ± 13.05 n = 7, ns p = 0.7023; mCherry 12h vs mCherry 48h, *** p < 0.001). In the POST, **(D)** c-Fos density was significantly higher in the mCherry 12h group compared to the mCherry 48h group, while chemogenetic manipulation did not significantly alter overall c-Fos density relative to respective controls (one-way ANOVA with Sidak’s multiple comparisons test: mCherry 12h 982.4 ± 89.39 n = 5 vs hM4Di 12h 855.8 ± 24.01 n = 7, ns p = 0.3459; mCherry 48h 731.2 ± 54.33 n = 5 vs hM3Dq 48h 787.5 ± 41.04 n = 8, ns p = 0.8628; mCherry 12h vs mCherry 48h, * p = 0.0239). Ensemble reactivation of learning-tagged neurons in the mPFC **(E)**. Chemogenetic inhibition significantly reduced the proportion of reactivated neurons, whereas chemogenetic stimulation induced c-Fos expression in nearly all tagged cells. Under control conditions, ensemble reactivation was significantly higher at 12h compared to 48h (one-way ANOVA with Sidak’s multiple comparisons test: mCherry 12h 42.47 ± 1.88 n = 6 vs hM4Di 12h 30.67 ± 1.32 n = 8, *** p < 0.001; mCherry 48h 30.61 ± 1.7 n = 6 vs hM3Dq 48h 93.97 ± 0.79 n = 6, *** p < 0.001; mCherry 12h vs mCherry 48h, *** p < 0.001). **(F)** Ensemble reactivation of learning-tagged neurons in the RSP. Chemogenetic inhibition significantly reduced the proportion of reactivated neurons compared to the mCherry 12h control group, whereas chemogenetic stimulation at 48h induced a strong increase in ensemble reactivation. Notably, ensemble reactivation was significantly higher in the mCherry 12h group compared to the mCherry 48h group (one-way ANOVA with Sidak’s multiple comparisons test: mCherry 12h 63.12 ± 1.94 n = 6 vs hM4Di 12h 32.56 ± 1.64 n = 7, *** p < 0.001; mCherry 48h 33.70 ± 1.69 n = 6 vs hM3Dq 48h 94.11 ± 0.62 n = 6, *** p < 0.001; mCherry 12h vs mCherry 48h, *** p < 0.001). **(G)** In the ORB, the proportion of reactivated neurons did not differ between the mCherry 12h and mCherry 48h control groups. Reactivation was significantly reduced in the hM4Di 12h group compared to its mCherry 12h control, whereas chemogenetic stimulation at 48h (hM3Dq 48h) resulted in higher ensemble reactivation (one-way ANOVA with Sidak’s multiple comparisons test: mCherry 12h 48.07 ± 2.18 n = 6 vs hM4Di 12h 29.50 ± 1.14 n = 8, *** p < 0.001; mCherry 48h 41.01 ± 3.94 n = 5 vs hM3Dq 48h 96.40 ± 0.78 n = 7, *** p < 0.001; mCherry 12h vs mCherry 48h, ns p = 0.1228). **(H)** Ensemble reactivation of learning-tagged neurons in the POST. The percentage of reactivated neurons was significantly higher in the mCherry 12h group compared to the mCherry 48h group. Chemogenetic inhibition significantly reduced ensemble reactivation relative to the mCherry 12h control, whereas chemogenetic stimulation at 48h (hM3Dq 48h) induced a strong increase in reactivation (one-way ANOVA with Sidak’s multiple comparisons test: mCherry 12h 71 ± 1.39 n = 5 vs hM4Di 12h 45.35 ± 1.12 n = 7, *** p < 0.001; mCherry 48h 45.91 ± 3.64 n = 5 vs hM3Dq 48h 94.97 ± 0.80 n = 8, *** p < 0.001; mCherry 12h vs mCherry 48h, *** p < 0.001). **(I)** Analysis of ensemble reactivation relative to chance level in control groups for the mPFC (mCherry 12h 1.518 ± 0.100, n = 6, ** p = 0.0035, df = 5, t = 5.175; mCherry 48h 1.059 ± 0.031, n = 6, ns p = 0.1177, df = 5, t = 1.888; one-sample t test. mCherry 12h vs mCherry 48h, ** p = 0.0014, df = 10, t = 4.375; unpaired t-test); **(J)** RSP (mCherry 12h 1.916 ± 0.138, n = 6, ** p = 0.0012, df = 5, t = 6.614; mCherry 48h 1.045 ± 0.061, n = 6, ns p = 0.5039, df = 5, t = 0.7198; one-sample t test. mCherry 12h vs mCherry 48h, *** p < 0.001, df = 10, t = 5.746; unpaired t-test); **(K)** ORB (mCherry 12h 1.187 ± 0.06, n = 6, ** p = 0.0029, df = 5, t = 5.398; mCherry 48h 1.156 ± 0.052, n = 5, * p = 0.0410, df = 4, t = 2.974; one-sample t test. mCherry 12h vs mCherry 48h, ns p = 0.0696, df = 9, t = 2.059; unpaired t-test); **(L)** POST (mCherry 12h 1.825 ± 0.161, n = 5, ** p = 0.0069, df = 4, t = 5.108; mCherry 48h 1.235 ± 0.068, n = 5, * p = 0.0264, df = 4, t = 3.435; one-sample t test. mCherry 12h vs mCherry 48h, ** p = 0.0098, df = 8, t = 3.367; unpaired t-test). **(M-P)** Relationship between ensemble reactivation and behavioral performance across brain regions during natural memory recall. Lines indicate linear regression fits (mPFC: mCherry 12h R^2^ = 0.0168, p = 0.8065; mCherry 48h R^2^ = 0.0158, p = 0.8120; RSP: mCherry 12h R^2^ = 0.6880, p = 0.0411; mCherry 48h R^2^ = 0.2770, p = 0.2834; ORB: mCherry 12h R^2^ = 0.5209, p = 0.1054; mCherry 48h R^2^ = 0.9729, p = 0.2740; POST: mCherry 12h R^2^ = 0.0001, p = 0.9825; mCherry 48h R^2^ = 0.0004, p = 0.9110). Data are presented as mean ± SEM.

### Simultaneous recordings of mPFC and RSP neural activity during memory recall

To further investigate the specific contribution of the mPFC and RSP during episodic-like memory recall, we analyzed changes in local field potential (LFP) signals simultaneously recorded from the two regions during the recall phase of the OPCRT in freely moving mice. LFP activity was recorded using two 8-channel electrode bundles and aligned to object exploration epochs; the location of individual recording sites was confirmed by post-hoc histological analysis (Figure **7A** and **7B**). Spectral analysis of mPFC activity during novel and familiar object exploration revealed a selective increase in low-frequency power during novel object interaction (Figure **7C** and **7D**). Focusing on the theta (5-10 Hz), low-gamma (30-55 Hz), and high-gamma (55-80 Hz) bands, we observed a significant increase in theta power during novel exploration (Figure **7E**). In the RSP, exploration of both novel and familiar objects was associated with a selective increase in theta-band activity; however, no significant differences were observed between the two conditions across the frequency bands considered (Figure **7F**). Interestingly, a modest increase in high-gamma power was observed during novel object exploration, although this effect was not statistically significant (Figure **7G** and **7H**). We next examined functional coupling between the mPFC and RSP by analyzing LFP coherence during novel and familiar object exploration (Figure **7I**). Coherence analysis provides a measure of frequency-specific synchronization between neural signals recorded from distinct brain regions. Coherence spectra obtained during novel and familiar exploration were largely overlapping, and no statistically significant differences were detected across the frequency range examined (Figure **7J**). Interestingly, for both conditions, coherence analysis produced a U-shaped profile, characterized by higher coherence values in the theta and high-gamma bands and lower coherence at intermediate frequencies. Finally, we assessed the directionality of mPFC-RSP interactions using Granger causality analysis during novel and familiar object exploration. Granger causality analysis assesses whether activity in one time series predicts future activity in another. When expressed in the frequency domain, this approach allows the investigation of directional interactions between brain areas in specific frequency bands. For both conditions, Granger causality was higher in the theta band for the RSP to mPFC direction compared to the mPFC to RSP direction, suggesting that RSP activity more strongly predicted mPFC activity during novel exploration (Figure **7K**-**7N**). By contrast, Granger causality in the low- and high-gamma bands was low and did not differ between directions. Notably, in the RSP to mPFC direction, Granger causality values were higher during familiar object exploration compared to novel object exploration, suggesting that the strength of directional interactions varies with recognition of the OPC association. This effect was specific to the theta frequency band and was not observed in the gamma ranges. Taken together, these results indicate that, during episodic-like memory recall, exploration of the novel object is associated with increased theta power in the mPFC, whereas no differences in theta or gamma power were detected in the RSP comparing novel and familiar exploration. In the theta band, directional interactions from RSP to mPFC were observed in both conditions and were more pronounced during familiar object exploration, suggesting that contextual input from the RSP partially drives prefrontal activity, particularly during familiarity recognition.

**Figure 7:**
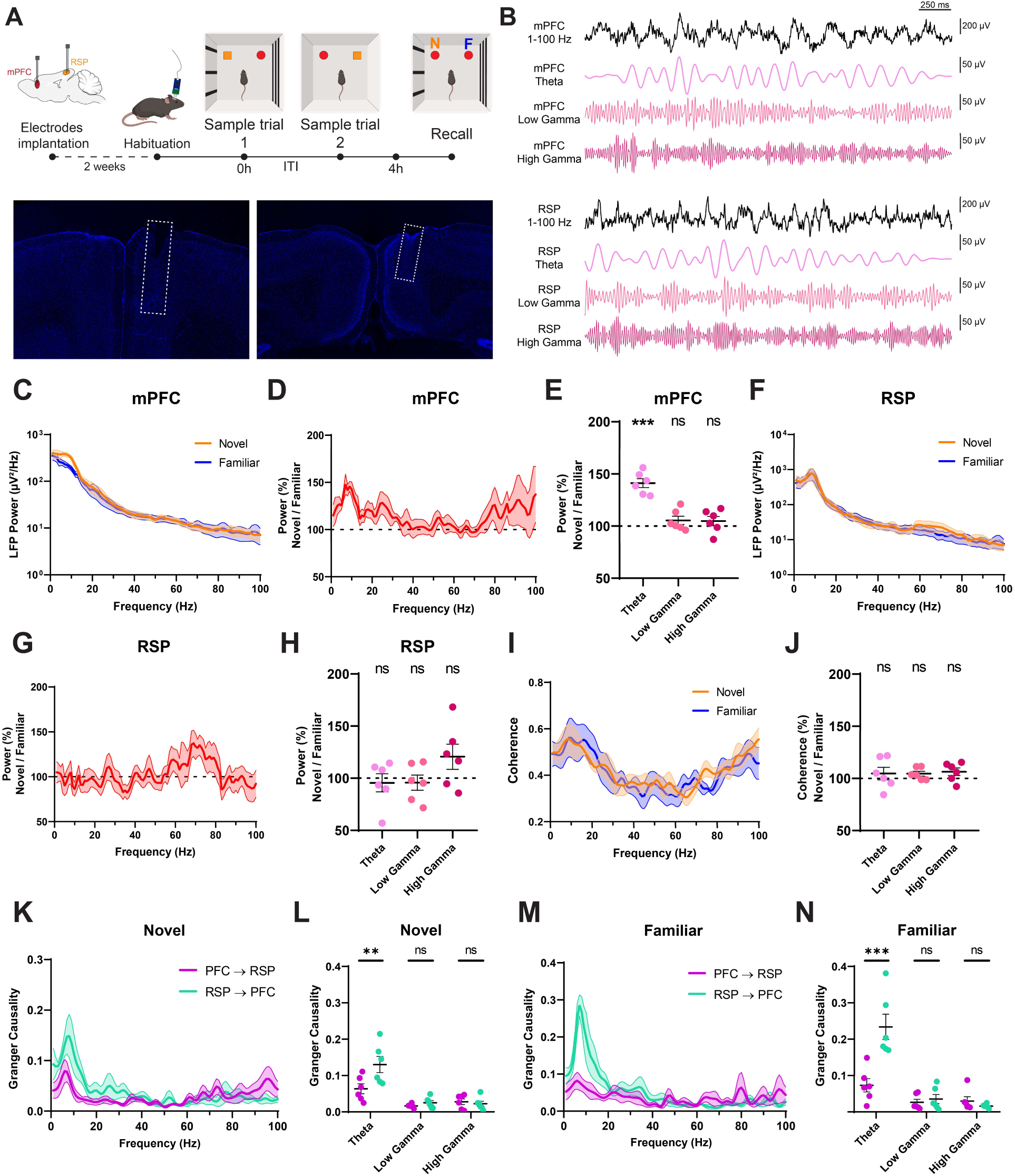
Simultaneous recordings of mPFC and RSP neural activity during memory recall. **(A)** In vivo simultaneous recordings from the medial prefrontal cortex (mPFC) and retrosplenial cortex (RSP) in freely moving mice during the recall phase of the object-place-context recognition task (OPCRT). Histological confirmation of electrode locations in the mPFC and RSP are shown below. **(B)** Representative examples of band-pass filtered LFP traces from both regions during object exploration. **(C)** mPFC power spectral density during novel and familiar object exploration epochs. **(D)** mPFC power expressed as the percentage of the novel exploration epochs relative to the familiar exploration epochs. **(E)** Band-averaged mPFC power in the theta (5-10 Hz), low-gamma (30-55 Hz), and high-gamma (55-80 Hz) frequency ranges, expressed as the percentage of the novel exploration epochs relative to the familiar exploration epochs. Theta-band power was significantly higher during novel compared to familiar exploration, whereas no significant differences were observed in the gamma bands (Theta 141.2 ± 4.290, n = 6, *** p = 0.0002, df = 5, t = 9.597; Low Gamma 105.4 ± 3.978, n = 6, ns p = 0.2317, df = 5, t = 1.361; High Gamma 104.9 ± 4.469, n = 6, ns p = 0.3272, df = 5, t = 1.086; one-sample t test). **(F)** RSP power spectral density during novel and familiar object exploration epochs. **(G)** mPFC-RSP power expressed as the percentage of the novel exploration epochs relative to the familiar exploration epochs. **(H)** Band-averaged RSP power during novel and familiar object exploration epochs. No significant differences were observed between conditions across the frequency bands examined (Theta 95.57 ± 8.686, n = 6, ns p = 0.6321, df = 5, t = 0.5094; Low Gamma 95.76 ± 7.286, n = 6, ns p = 0.5863, df = 5, t = 0.5812; High Gamma 120.6 ± 12.09, n = 6, ns p = 0.1486, df = 5, t = 1.707; one-sample t test). **(I)** Coherence between mPFC and RSP during novel and familiar object exploration epochs. Coherence spectra were computed across the 1-100 Hz range and exhibited comparable profiles between conditions. **(J)** mPFC-RSP coherence expressed as the percentage of the novel exploration epochs relative to the familiar exploration epochs. No significant differences were observed between conditions across the frequency bands considered (Theta 104.7 ± 5967, n = 6, ns p = 0.4675, df = 5, t = 0.7859; Low Gamma 104.6 ± 2.347, n = 6, ns p = 0.1098, df = 5, t = 0.1.942; High Gamma 106.4 ± 3.635, n = 6, ns p = 0.1366, df = 5, t = 1.772; one-sample t test). **(K)** Granger causality spectra during novel object exploration epochs for both directions of interaction between RSP and mPFC. **(L)** During novel exploration, Granger causality in the RSP to mPFC direction was significantly higher than in the mPFC to RSP direction (two-way ANOVA with Sidak’s multiple comparisons test: PFC to RSP Theta 0.064 ± 0.014 n = 6 vs RSP to PFC Theta 0.130 ± 0.022 n = 6, ** p = 0.0011; PFC to RSP Low Gamma 0.017 ± 0.002 n = 6 vs RSP to PFC Low Gamma 0.025 ± 0.006 n = 6, ns p = 0.9400; PFC to RSP High Gamma 0.028 ± 0.008 n = 6 vs RSP to PFC High Gamma 0.022 ± 0.007 n = 6, ns p = 0.9836). **(M)** Granger causality spectra during familiar object exploration epochs for both directions of interaction between RSP and mPFC. **(N)** During familiar exploration, Granger causality in the RSP to mPFC direction was significantly higher than in the mPFC to RSP direction (two-way ANOVA with Sidak’s multiple comparisons test: PFC to RSP Theta 0.073 ± 0.018 n = 6 vs RSP to PFC Theta 0.234 ± 0.035 n = 6, *** p < 0.001; PFC to RSP Low Gamma 0.026 ± 0.008 n = 6 vs RSP to PFC Low Gamma 0.035 ± 0.013 n = 6, ns p = 0.9795; PFC to RSP High Gamma 0.030 ± 0.012 n = 6 vs RSP to PFC High Gamma 0.016 ± 0.002 n = 6, ns p = 0.9273). Scale bars are shown in the images. Data are presented as mean ± SEM.

To further investigate the activity of the examined brain regions during the recall phase of the OPCRT, we analyzed single neurons identified by single-unit isolation during interactions with novel and familiar object configurations (Figure **8A** and **8B**). Following criteria established in previous studies^49,50^, all identified units were classified as putative regular-spiking (RS) excitatory neurons, as they exhibited baseline firing rates below 10 Hz and spike widths longer than 0.3 ms (Figure **8C**). mPFC and RSP neurons exhibited distinct temporal dynamics during object exploration, with firing rate modulation dependent on object configuration (Figures **8D** and **8E**). In the mPFC (Figure **8D**), neuronal activity displayed a clear modulation by object novelty. Specifically, a transient increase in firing rate was observed during the pre-exploration phase selectively in the novel condition, while no comparable peak was evident for the familiar configuration. During the exploration phase, both novel and familiar conditions elicited an increase in activity; however, the response to novel objects was consistently larger in magnitude. In the RSP (Figure **8E**), a partially distinct profile emerged. No evident modulation was observed during the pre-exploration phase for either condition. In contrast, during exploration, both novel and familiar configurations elicited a sharp increase in firing rate, similar to mPFC. However, as in mPFC, the response to the novel configuration was stronger, suggesting an enhanced recruitment of RSP neurons during the processing of novelty. To quantitatively assess these differences, we compared firing rates between novel and familiar conditions across epochs. During the pre-exploration phase (Figure **8F**), mPFC activity was significantly higher in the novel configuration compared to the familiar one, whereas no significant difference was observed in the RSP. During the exploration phase (Figure **8G**), firing rates were significantly higher for novel compared to familiar configurations in both mPFC and RSP. Together, these results indicate that novelty-related modulation emerges earlier in mPFC activity, already during pre-exploration, while RSP engagement appears more tightly linked to the exploration phase.

**Figure 8:**
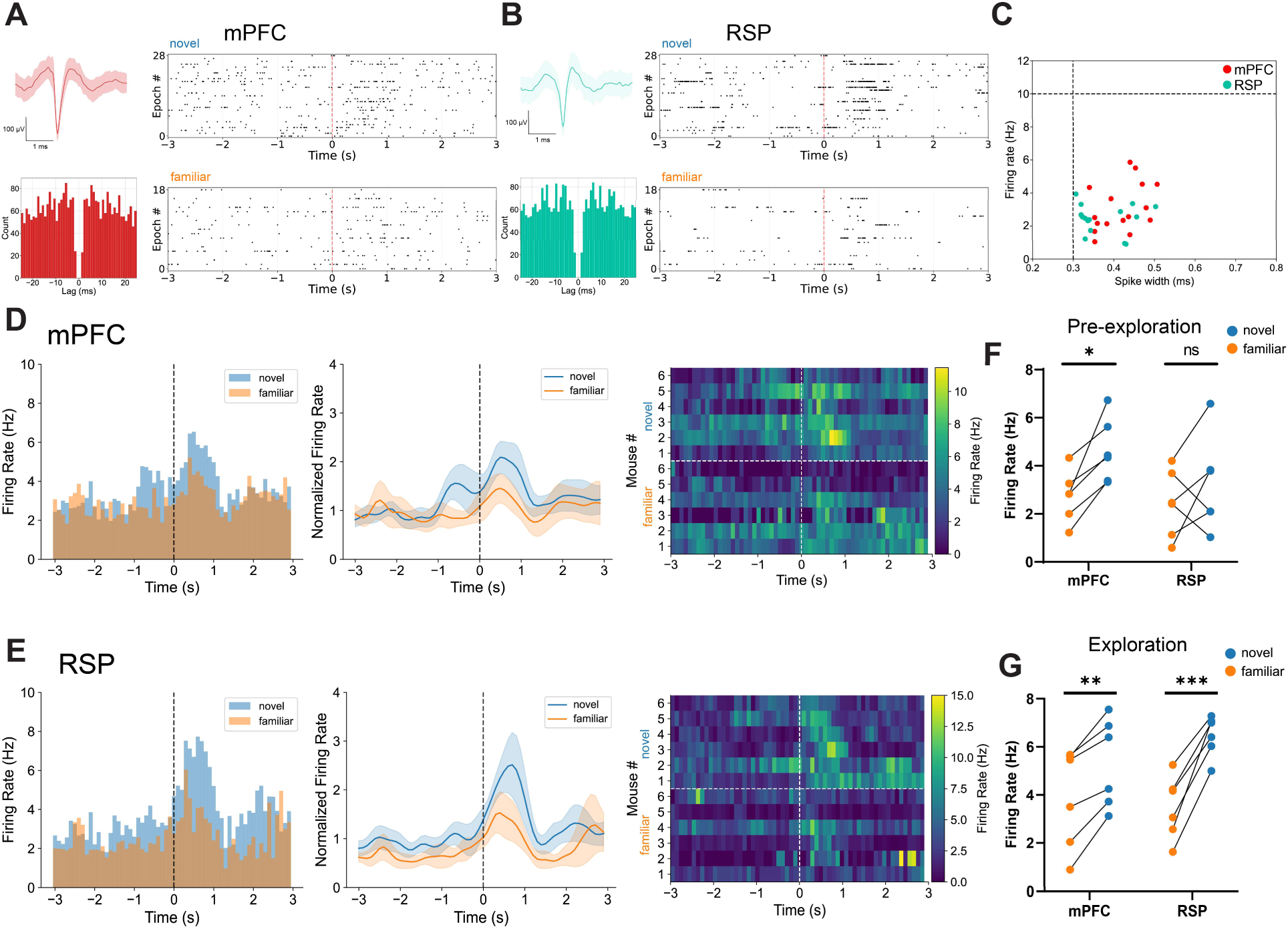
Single-unit recordings in mPFC and RSP during OPC memory recall. **(A-B)** Representative single units recorded in mPFC **(A)** and RSP **(B)**, showing mean spike waveform, autocorrelogram, and spike raster plots during exploration epochs of the novel configuration (top) and familiar configuration (bottom). **(C)** Spike waveform width (ms) and firing rate (Hz) across units recorded in mPFC (n = 17 units) and RSP (n = 16 units) from 6 mice. All units exhibit spike widths > 0.3 ms and firing rates < 10 Hz and are classified as putative regular-spiking (RS) neurons. **(D-E)** Peri-stimulus time histograms (PSTHs; 0.1 s bins) aligned to object exploration for mPFC **(D)** and RSP **(E)**. Left panels show raw firing rate responses during novel and familiar exploration epochs. Middle panels show PSTHs normalized to baseline activity. Right panels display heatmaps of mean activity across mice for each exploration condition, illustrating the temporal dynamics of neural responses during novel and familiar exploration. **(F)** Comparison of population activity between novel and familiar exploration in mPFC and RSP during the pre-exploration period (−1 to 0 s). The mPFC shows a significant increase in activity prior to novel exploration relative to familiar conditions (two-way RM ANOVA with Sidak’s multiple comparisons test: mPFC Novel 4.639 ± 0.542 n = 6 vs mPFC Familiar 2.747 ± 0.435 n = 6, * p = 0.0292; RSP Novel 3.239 ± 0.802 n = 6 vs RSP Familiar 2.415 ± 0.571 n = 6, ns p = 0.4053). **(G)** Comparison of population activity between novel and familiar exploration in mPFC and RSP during the exploration period (0 to +1 s). Both regions exhibit strong activation following novel exploration (two-way RM ANOVA with Sidak’s multiple comparisons test: mPFC Novel 5.319 ± 0.754 n = 6 vs mPFC Familiar 3.859 ± 0.839 n = 6, ** p = 0.0016; RSP Novel 6.463 ± 0.350 n = 6 vs RSP Familiar 3.486 ± 0.535 n = 6, *** p < 0.001). Data are presented as mean ± SEM.

## Discussion

Using a brain-wide mapping approach, we showed that episodic-like memory retrieval recruits a fronto-posterior medial network of brain regions beyond the classical involvement of the medial temporal lobe system. This supports the idea that memory traces are encoded across distributed neuronal populations, in line with early theoretical proposals by Richard Semon^51^. Chemogenetic manipulation of learning-tagged ensembles within this network demonstrated that, although all the examined regions were necessary for successful memory expression, only the RSP satisfied both sufficiency and necessity criteria, suggesting that it contains an engram population for this memory. In vivo recordings showed that mPFC theta power increased during novel exploration, along with a transient increase in firing rate before and during exploration of the novel configuration. In addition, RSP neurons exhibited a sharp increase in firing following exploration of the novel configuration.

To characterize the patterns of neuronal activity specifically associated with episodic-like memory, we mapped c-Fos expression across the entire brain following behavioral testing. Analysis of the regional contributions to the principal components driving condition separation indicated an important involvement of isocortical areas across the first three components. Cortical activation has been observed during spatial and learning tasks across parietal, occipital, and cingulate areas, and the isocortex has been proposed to function as a multimodal associative system involved in constructing spatial representations of the environment^52–54^. In the comparison of the OPCRT group with the HC condition, task PLS analysis confirmed a widespread increase in c-Fos expression across cortical and subcortical regions, reflecting the global engagement induced by exploration. When OPCRT was compared with the CNTX control, the number of significantly activated regions was reduced. Notably, primary sensory cortices remained significantly engaged, suggesting that object interaction drives a strong activation of these regions. In addition, associative cortical regions, including frontal areas and the RSP, showed increased activation, together with selective recruitment of hippocampal and parahippocampal regions such as CA3 and the POST. This pattern suggests that, beyond sensorimotor processing, OPCRT engages higher-order associative circuits involved in integrating object-related information with contextual features^55,56^. Finally, comparison with the OCT condition, which controls for both spatial navigation and object interaction, resulted in a marked restriction of the activated network. The substantial reduction in significantly activated regions indicates that most areas identified in the previous contrasts are primarily associated with exploration, sensory processing, and object interaction rather than with memory retrieval. Although the OCT condition engages memory retrieval processes, the nature of the retrieved information differs fundamentally from that required for episodic-like memory. In this condition, during the recall phase no novel OPC configuration or sensory mismatch is introduced, and all stimulus features remain identical to those encountered during part of the learning phase. In the absence of a mismatch between expected and experienced stimulus configurations, there is no requirement to retrieve and compare a stored OPC association. Consequently, novelty-driven exploration and prediction-error signals that normally trigger episodic-like retrieval are not engaged^57,58^. Under these conditions, retrieval of the OPC association, even if stored, is neither required nor behaviorally expressed, and task performance can be supported by recognition of individual stimulus elements without accessing their associative configuration within a specific episode. In comparison with the OCT control, only a limited subset of regions remained selectively activated during OPCRT, including the mPFC, ORB, RSP and POST. This analysis reveals a hierarchical organization of recruited brain regions that reflects the increasing cognitive demands of the control conditions. The progressive restriction of identified activated areas suggests that the increasing cognitive load of the tasks effectively filter out regions primarily involved in sensory processing and object exploration, isolating a core network of regions specifically implicated in OPC memory retrieval.

Beyond changes in regional activation, memory retrieval is expected to engage coordinated interactions across multiple brain systems. We therefore examined the co-variation of c-Fos expression across areas, an approach that has been applied to study functional networks^12,59^. Episodic-like memory retrieval was associated with a marked reorganization of large-scale functional interactions, suggesting a more efficient integration of information across brain regions. Notably, this reconfiguration was accompanied by an increase in small-world network properties, characterized by a balance between enhanced global communication and preserved local clustering^60^. Such a topology is thought to optimize information transfer while minimizing processing costs, supporting efficient communication between distributed neural assemblies^61,62^. Analysis of topological roles suggests that the POST and RSP occupy a central position in mediating communication between distinct functional modules, reinforcing the idea of an involvement of posterior medial structures in the network engaged during episodic-like memory retrieval^63^.

While brain-wide analyses of neuronal activation during OPCRT recall identified a core subset of selectively engaged brain regions, activity alone was insufficient to define the contribution or functional role of individual neuronal populations. By combining TRAP-based activity-dependent tagging with chemogenetic manipulation, we directly tested whether learning-activated neuronal ensembles are necessary and/or sufficient for memory expression. This approach has previously been applied to characterize learning-activated neurons in the LEC, providing causal evidence for the existence of a neuronal subpopulation with properties of engram cells for OPC memory recall^31,32^. Across all regions examined, chemogenetic inhibition of learning-tagged neuronal populations consistently impaired OPCRT performance, suggesting that these ensembles are not only active during recall but are necessary for successful memory expression. At the behavioral level, this impairment manifested as a loss of preference for the object presented in the novel configuration, indicating a failure to appropriately retrieve or utilize OPC associations. Importantly, although the behavioral outcome was similar across regions, the underlying mechanisms leading to this deficit are likely to be region specific. For instance, inhibition of learning-tagged neurons in the mPFC may disrupt higher-order processes required for the integration and selection of task-relevant mnemonic information, impairing the expression of the appropriate behavioral response^64,65^. In the ORB, the observed deficit may reflect a failure to assign behavioral significance of competing object configurations, consistent with a role in representing task states or expected outcomes^66,67^. The disruption of activity of learning-tagged neurons in the POST may impair the integration of spatial information necessary to anchor object representations within a context, preventing discrimination between familiar and novel configurations^68,69^. In contrast, reactivation of learning-tagged neurons in the RSP was sufficient to restore memory expression when natural cues were no longer effective, thus inhibition of this population is likely to directly interfere with reactivation of the memory trace itself. This suggests that learning-tagged RSP neurons possess functional properties of engram cells. Consistent with this interpretation, the anatomical position and connectivity of the RSP place it in a unique position to act as an interface between regions involved in sensorimotor and spatial processing and those engaged in mnemonic and associative functions^44,70,71^. Analysis of ensemble reactivation suggests that learning-tagged populations in the ACA and ORB may be recruited in shaping the coordination of network activity or motivational state during recall, rather than supporting the selective reactivation of the memory trace. The PL and the RSP exhibited a significant relationship between the proportion of ensemble reactivation and behavioral performance. In both areas, higher levels of reactivation of learning-tagged neurons were associated with high discrimination performance, suggesting a direct role in driving memory-guided behavior.

However, these approaches provide limited temporal resolution to characterize the dynamics of neural activity in individual regions. To investigate how neural activity changes during exploration of novel and familiar configurations, we conducted electrophysiological recordings in freely moving animals during the recall phase of the OPCRT. We focused on local field potential (LFP) signals, which reflect the integrated excitatory and inhibitory synaptic activity of local neuronal populations within a radius of a few hundred micrometers around the recording electrode^72,73^. Consistent with previous findings, power spectral analysis revealed that LFP activity in both the mPFC and RSP was dominated by theta-frequency components^74^. Comparing novel and familiar exploration epochs, we observed that novel object exploration was associated with increased theta power in the mPFC. Because phase synchronization has been proposed as a mechanism underlying coordination between brain regions during behavior and cognition^75,76^, we quantified functional coupling between the mPFC and RSP using coherence and Granger causality analyses. Coherence analysis indicated that novel and familiar exploration epochs did not differ in the degree of phase consistency between the mPFC and RSP. The absence of condition-dependent changes in coherence may also reflect the similarity of exploratory behavior between novel and familiar conditions, potentially masking more subtle differences related to memory recall or novelty detection. Granger causality analysis revealed that directional interactions between mPFC and RSP were frequency-specific and predominantly expressed in the theta band. In both novel and familiar exploration conditions, theta-band Granger causality was consistently higher in the RSP to mPFC direction than in the opposite. Given the possible role of the RSP in integrating contextual information, this directional coupling may reflect the transmission of previous contextual representations to prefrontal circuits, facilitating the access to previously learned OPC associations.

At the level of single-neuron activity, our findings suggest a temporally structured contribution of mPFC and RSP to novelty processing, characterized by earlier novelty-related modulation in mPFC and a novelty-dependent engagement of both regions during exploration. The early increase in mPFC activity prior to object interaction is consistent with evidence from cognitive tasks showing that mPFC unit activity can predict future choice behavior^77^. In fact, prefrontal circuits have been proposed to encode expectations about upcoming stimuli and to signal deviations from predicted outcomes^78,79^. The enhanced mPFC response observed immediately before exploration of the novel configuration may reflect the attribution of behavioral salience to the unexpected change in object arrangement, promoting investigation of the novel object. Previous work has shown that RSP neurons are sensitive to the egocentric distance and direction of both spatial locations and discrete objects in the environment^44^. In addition, neurons within the RSP respond preferentially to objects that carry strong contextual significance, including objects defined by their expected spatial location or by learned associations^80^. In the OPCRT, when the familiar object arrangement was altered, the animal was required to update previously acquired associations between object identity, position and context. Under these conditions, the increased RSP activity may reflect the comparison between stored contextual representations and incoming sensory information.

Based on the activation patterns observed in the brain-wide c-Fos mapping and the region-specific functional dissection obtained through chemogenetic manipulation, we hypothesize that a network composed of a frontal and posterior medial region is activated during OPC memory recall (Figure **9A**). Within this network, the RSP appears as a critical node containing neuronal populations with characteristics of engram cells for OPC memory. In addition, previous evidence showed that episodic-like memory is associated with synaptic plasticity changes in the LEC, and that this region contains neuronal populations that are both necessary and sufficient for OPC memory recall^31^. These results support a model in which episodic-like memory in mice is sustained by a distributed network, with the RSP and LEC representing brain regions fundamental for successful memory retrieval. Other regions within this network appear to contribute to memory recall by supporting the correct behavioral expression of the memory rather than directly storing its content. Based on the major anatomical connections between these brain regions^46^, the LEC emerges as a key area linking the frontal and posterior medial components of the network. Notably, information from the RSP and POST is likely conveyed to the LEC through strong connections with the MEC; however, this subdivision appears to be less directly involved in OPC memory recall^31^. However, the relationship between the activated neuronal ensembles in the LEC and RSP remains to be clarified, particularly whether they operate as coordinated components of a distributed engram or contribute complementary information through parallel yet interacting processes, or whether activity in one region drives and shapes the activation of the other. Overall, our findings indicate a complex organization of episodic-like memory, composed of multiple interconnected brain regions that are necessary for successful memory recall. While only the RSP showed neuronal populations with properties consistent with engram cells, the other identified areas critically contribute to memory expression, although the specific contribution of each region remains to be determined.

**Figure 9:**
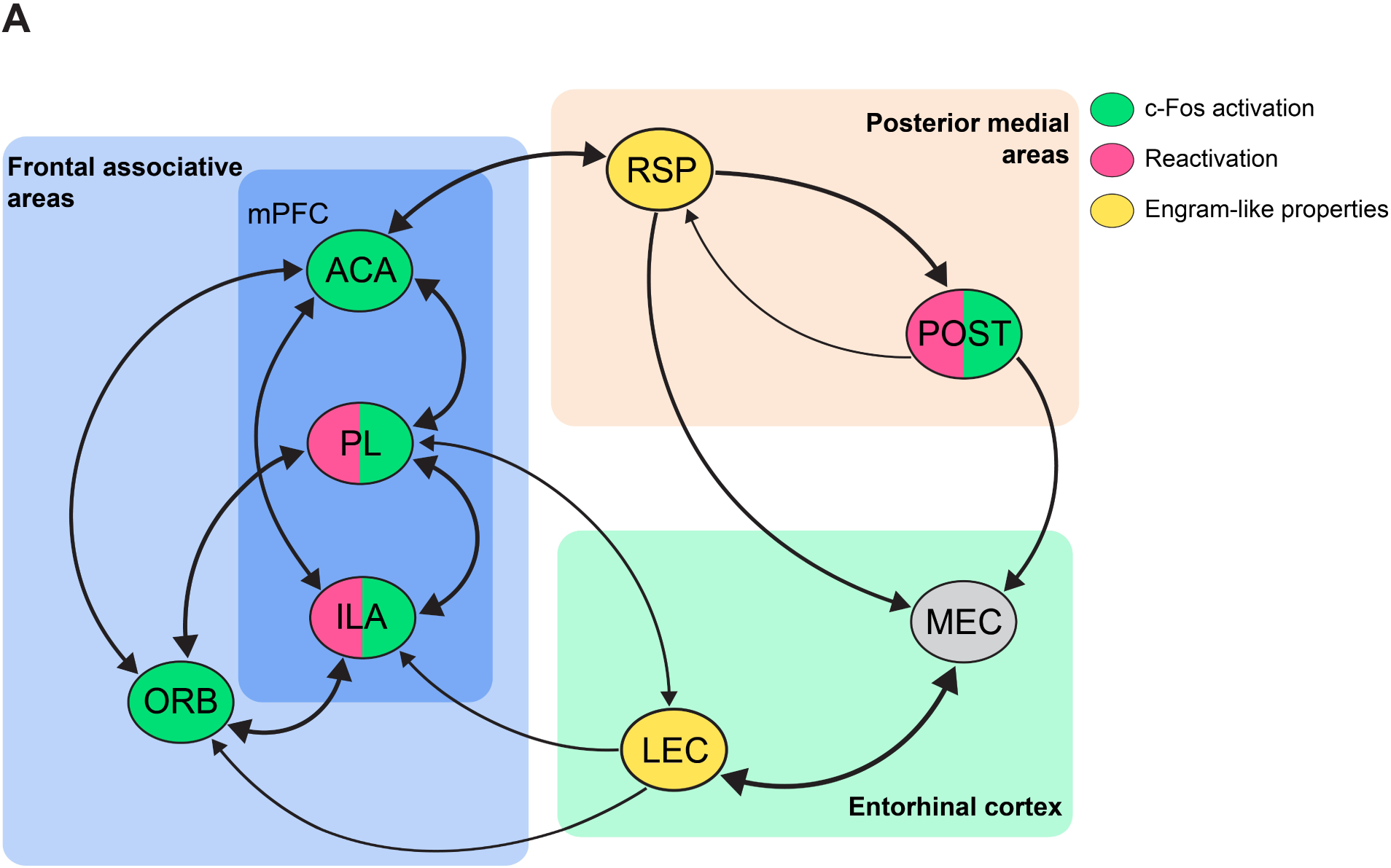
Proposed distributed network for episodic-like memory. **(A)** The schematic illustrates a distributed network organized into three major components: frontal associative areas, posterior medial areas, and the entorhinal cortex. Nodes represent brain regions identified as part of the OPC memory network based on the present study and previous work. Green nodes indicate brain regions that showed increased c-Fos activation during OPC memory recall at 12h compared to 48h. Red nodes indicate regions in which learning-tagged neuronal ensembles exhibited significantly higher reactivation at 12h relative to 48h. Yellow nodes highlight brain regions containing learning-tagged neuronal populations with properties of engram cells for OPC memory, as demonstrated by both necessity and sufficiency for memory recall. The MEC is included based on its anatomical position linking posterior medial areas with the LEC. Edges represent major direct anatomical connections between regions, with edge width proportional to connection density.

## Materials and methods

### Animals

All experimental procedures involving animals followed the guidelines defined by the European legislation (Directive 2010/63/EU), and the Italian Legislation (LD no. 26/2014). The Organism Responsible for Animal Welfare (OPBA) of the National Research Council of Italy (CNR) Institute of Neuroscience in Pisa and the Italian Ministry of Health approved the study protocol (authorization n. 16/2022-PR). Mice were housed in conventional cages (365 x 207 x140 mm, 2-3 animals per cage) with nesting material on a 12-h light/dark cycle with food and water available ad libitum. For brain-wide c-Fos mapping, experiments were performed in male wild-type C57BL/6J mice (Charles River, Italy) at 3 months of age. For chemogenetic manipulation and electrophysiology experiments, both male and female wild-type C57BL/6J mice at 3 months of age were used. All behavioral procedures were performed during the light phase, and mice were randomly assigned to experimental groups. To control for order and cage effects, each cage contained a mixture of mice from the experimental and control groups. The number of animals used in each experiment is provided in the figure legends.

### AAV vectors and stereotaxic injections

The following AAV vectors were used for the chemogenetic manipulation: AAV5-Fos::CreER^T2^ (titer 1.2x10^13^), AAV5-hSyn::DIO-hM3Dq-mCherry, AAV5-hSyn::DIO-hM4Di-mCherry and AAV5-hSyn::DIO-mCherry (titers: 5.0-6.0x10^12^). For stereotaxic injections, 2-month-old mice were deeply anesthetized using an intraperitoneal injection of Zoletil 100 (zolazepam and tiletamine, 1:1, 40 mg/kg; Laboratoire Virbac) and Xilor (xylazine 2%, 10 mg/kg; Bio98). Lidocaine (2%) was topically applied to the skull to provide local analgesia. A bilateral craniotomy was performed at the stereotaxic coordinates targeting: mPFC (AP +1.8 mm, ML ± 0.2 mm, DV -1.55 mm), RSP (AP -2.5 mm, ML ± 0.2 mm, DV -0.7 mm), ORB (AP +2.6 mm, ML ± 1.0 mm, DV -1.95 mm), and POST (AP -4.2 mm, ML ± 2.1 mm, DV -1.9 mm). Dorso-ventral coordinates are reported relative to the brain surface. For all regions, a viral mixture of AAV5-FosCreER^T2^ and Cre-dependent AAV (ratio 1:500, AAV5-FosCreER^T2^ at a final titer of 2.43x10^10^) was injected. Injections were performed bilaterally using a glass micropipette connected to a microinjection system at a constant flow rate of 0.1 μL/min. Injection volumes were region-specific: 300 nL were delivered into the mPFC and ORB, whereas 250 nL were injected into the RSP and POST. Following each injection, the pipette was left in place for an additional 5 min to allow diffusion of the viral solution before slow withdrawal. After surgery, animals were returned to their home cages and allowed to recover for 3 weeks before the start of behavioral experiments.

### Behavioral tasks

The test environment was composed of two square boxes (length 40cm, width 40cm, height 40cm) with different visual cues on the walls to provide distinct contexts. The wall and floor of the environment were cleaned with 30% alcohol before each trial. The objects were household items of approximately the same size as the mouse and varying in color, shape, and texture. To avoid odor cues, new identical copies of each object were used for each trial, and objects were cleaned with 30% alcohol after each trial.

Before the behavioral tests, mice were habituated to the experimenter and room by extensive handling for two weeks. For the Novel Object-Place-Context Recognition Task (OPCRT), mice were allowed to explore the first context for 5 minutes before the beginning of the sample phases. Subsequently, mice were presented with two distinct novel objects in Context A during the first sample trial. After this trial, the mice were removed from the box and placed in a holding cage for a 1-minute inter-trial interval (ITI) while the box was cleaned. In the second sample trial, the same pair of objects was presented in Context B, but their positions were reversed relative to the first trial. During the test trial, mice were exposed to a pair of identical objects from the sample trials in the same context and positions as in the first sample trial. Both the sample phases and the test phase were conducted for a duration of 10 minutes each. Animals were tested after a retention interval of 4h, 12h, or 48h, depending on the experimental group, as indicated in the figures.

For the context-exposure group (CNTX), mice were exposed to the same two contexts used in the OPCRT without the presence of objects, following the same context sequence, exposure duration, and retention interval as in the OPCRT.

For the object-context task (OCT), mice were exposed to the same two contexts used in the OPCRT while maintaining the same pair of objects in the same positions across all sample trials, following the same context sequence, exposure duration, and retention interval as in the OPCRT.

In all the experiments, mice were judged to be exploring an object when it was in proximity with their nose directed towards it. Exploration time was not counted when the mouse’s nose was directed away from the object. To ensure reliability, the same separate observer re-scored all the videos in a blind fashion for each task, and these scores consistently differed by less than 10% from those of the experimenter. For each task, observation scores were converted into discrimination indices (discrimination index = (time at novel − time at familiar) / (time at novel + time at familiar)) to evaluate the extent to which mice explored novel versus familiar objects. The scoring of the videos was performed using Chronotate^81^, a software designed for precise manual analysis of behavior during experimental trials. The marker output files generated by Chronotate were processed using a custom Python script, which allowed for the quantification of the total exploration time as well as the total number of interactions with both the novel and familiar objects. Behavioral videos were recorded using an AUKEY 1080p full HD webcam and were subsequently analyzed offline. Different body regions of the mouse, namely the nose, two ears, back, middle portion, and tip of the tail, were labeled using the open-source tool DeepLabCut^82^ for markerless pose estimation. This animal tracking enabled the automated calculation of various motor parameters, facilitating a more detailed analysis of the animal’s behavior.

### Pharmacological treatment

4OH-TAM (H6278, Sigma-Aldrich) was prepared for injection by dissolving it in saline. A stock solution of 50 mg/mL 4OH-TAM in DMSO (D8418, Sigma-Aldrich) was initially prepared and stored at -20°C. On the day of the experiment, the final working solution of 2.5 mg/mL 4OH-TAM was achieved in two stages: first, by diluting the stock 1:10 in saline that contained 2% Tween80 (P1754, Sigma-Aldrich), and then adding a volume of saline. This final solution consisted of 2.5 mg/mL 4OH-TAM, 5% DMSO, and 1% Tween80 in saline. Mice were given an intraperitoneal injection of 4OH-TAM at a dose of 25 mg per kg body weight, 4 hours prior to the sample trials.

Clozapine N-oxide hydrochloride (CNO; cat. no. 34233-7, Merck) was dissolved in sterile saline for injection. For the behavioral experiments, mice were administered 3 mg per kg (i.p.) of CNO 30 minutes before each test phase.

### Immunohistochemistry

Mice were deeply anesthetized with urethane (Merck, 20% solution, 0.1 mL/100 g body weight) and perfused intracardially with PBS (pH 7.4), followed by 4% paraformaldehyde (PFA) in PBS (pH 7.4). The brains were carefully removed, post-fixed overnight in 4% PFA (w/v), and transferred to a 30% sucrose solution (w/v) in PBS for dehydration and cryoprotection. Coronal sections, 50 µm thick, were prepared using a Leica freezing microtome, spanning from the main olfactory bulbs to the cerebellum, and the free-floating slices were subsequently processed for immunofluorescence analysis. For the brain-wide c-Fos mapping, one out of every 3 sections was collected for further processing leading to a sampling of one slice every 150 μm. For each animal, slices were assigned a unique ID and pooled in a 24-well plate for free-floating staining. Each well contained 5-6 sections that sampled the brain at an equally spaced position in the anterior-posterior axis. For the other experiments, the appropriate brain slices were manually selected.

All sections were incubated for 2h at room temperature in a blocking solution containing 5% BSA (w/v) and 0.5% Triton X-100 (v/v) in PBS. For brain-wide c-Fos mapping, sections were incubated overnight at 4°C with a monoclonal anti-c-Fos antibody (cat. no. 226 008, Synaptic Systems) diluted 1:1000 in PBS containing 1% BSA (w/v) and 0.1% Triton X-100 (v/v). Sections were then washed three times for 5 min in PBS and incubated for 2 h at room temperature with an Alexa Fluor 488-conjugated anti-rabbit secondary antibody (cat. no. 711-545-152, Jackson ImmunoResearch) diluted 1:500 in the same solution. For the other experiments, sections were incubated overnight at 4°C with the anti-c-Fos antibody together with a monoclonal anti-mCherry antibody (cat. no. M11217, ThermoFisher Scientific), both diluted 1:1000 in PBS containing 1% BSA and 0.1% Triton X-100. Corresponding Alexa Fluor 488-conjugated anti-rabbit and Alexa Fluor 568-conjugated anti-rat secondary antibodies (cat. nos. 711-545-152, Jackson ImmunoResearch; A11077, ThermoFisher Scientific) were applied for 2 h at room temperature at a dilution of 1:500. Finally, sections were washed three times in PBS, mounted onto glass slides, air-dried, and coverslipped using Fluoromount™ aqueous mounting medium (cat. no. F4680, Merck).

### Image acquisition

Fluorescence images were acquired using a Leica Stellaris 8 confocal microscope equipped with 10x/0.3 NA dry and 20x/0.75 NA dry objectives. For brain-wide c-Fos mapping, whole coronal sections were imaged using the 10x objective at a scan speed of 600 Hz and a resolution of 2048 x 2048 pixels, acquiring the optical plane exhibiting the maximal c-Fos signal. For the other experiments, tiled acquisitions spanning 4-6 z-planes were performed using the 10x or 20x objectives at a scan speed of 600 Hz and a resolution of 1024 x 1024 pixels.

### Image processing and registration

Image registration to the Allen Brain Atlas CCFv3-2017 was performed as previously described^37^. For each brain, images were ordered along the posterior-anterior axis according to their unique slice ID. All images were visually inspected and, when necessary, mirrored to ensure consistent left-right hemisphere orientation across sections. To restrict the analysis of c-Fos^+^ cell density to image regions containing brain tissue, binary masks were generated for each image using an Ilastik-trained machine learning model. All masks were manually inspected and, when required, corrected using a custom MATLAB script. Alignment to the Allen Brain Atlas was performed in three steps. First, experimental images were matched to the corresponding atlas planes using QuickNII v2.2. Alignment was then refined in VisuAlign v0.9 by manually applying local transformations based on anatomical landmarks and slice-specific distortions. Finally, each image pixel was mapped to its corresponding voxel in the reference atlas. Brain-wide heatmaps of c-Fos density fold-change were generated in Python using the BraiAnalyse library^83^.

### Cell counting

For brain-wide c-Fos mapping, c-Fos^+^ cells were detected using a custom Python pipeline based on StarDist^84^, a deep learning-based method for cell and nuclear segmentation. The raw segmentation output was subsequently refined by applying semi-automated thresholds on fluorescence intensity and object area to exclude background signal and segmentation artifacts. Threshold values were manually adjusted on a per-image basis when required to account for variations in staining quality and background fluorescence. All adjustments were visually inspected to ensure accurate identification of c-Fos^+^ while minimizing technical artifacts.

For the other experiments, regions of interest (ROIs) were manually outlined in ImageJ based on anatomical boundaries defined using the mouse brain atlas (Paxinos and Franklin, The Mouse Brain in Stereotaxic Coordinates, 2004). Detection of c-Fos^+^ cells within these ROIs was performed using the same custom Python pipeline described for brain-wide analysis. c-Fos^+^ cell density (c-Fos^+^/mm2) was calculated and averaged across 4-5 sections per animal. The same procedure was applied to quantify DAPI^+^ cells. For the mCherry^+^ cell detection, cell counting was manually performed in ImageJ and the mCherry^+^ cell density (mCherry^+^/mm2) was averaged over 4-5 sections per animal. The number of c-Fos^+^/mCherry^+^ cells per section was manually counted in ImageJ. Z-planes were examined to ensure accurate co-localization of the c-Fos and mCherry signals within the same cell. The percentage of reactivation within the tagged neuronal population was calculated as ((c-Fos^+^ mCherry^+^)/(mCherry^+^)) x 100. To assess the level of reactivation, the proportion of overlap with DAPI^+^was calculated as (c-Fos^+^mCherry^+^)/DAPI^+^. The expected chance level of overlap was determined as (c-Fos^+^/DAPI^+^) x (mCherry^+^/DAPI^+^), and this calculation was performed for each slice. Overlap/chance was then computed by dividing the overlap/DAPI value by the chance value for each slice. Overlap/chance was averaged across all slices for each mouse, generating a single average value used for statistical analysis.

### In vivo electrophysiology

Custom electrode assemblies were prepared by inserting Formvar-insulated nichrome wires (50.8 µm diameter, A-M systems, Cat. #762000) into the channel holes of a gold-plated electronic interface board (EIB-16, Neuralynx), forming two electrode bundles of eight electrodes each, which were secured with gold pins (Neuralynx). Ground/reference channels were connected to a silver wire soldered to a stainless-steel screw. For chronic implantation, 3-month-old C57BL/6J mice were deeply anesthetized with isoflurane (3% for induction, 1.5% for maintenance) and positioned in a stereotaxic frame. The scalp was disinfected with 70% ethanol followed by betadine solution before incision. Craniotomies were performed stereotaxically above the mPFC and RSP using the same coordinates described for viral injections, and an additional hole was drilled above the cerebellum for ground/reference placement. Electrode bundles were slowly lowered into the target regions to the appropriate dorsoventral coordinates. The stainless-steel screw connected to the reference/ground channel was implanted over the cerebellum. The EIB and ground screw were secured to the skull using dental cement (Super-Bond C&B) and further reinforced with acrylic dental cement (Paladur). Following surgery, mice were allowed to recover from anesthesia at 37°C before they returned to their home cages.

Recordings were performed 15 days after the surgery using an Open Ephys acquisition board and the Open Ephys GUI data acquisition software. Briefly, signals from the EIB-16 were amplified, filtered, and digitized using a 16-channel Intan RHD headstage and acquired via the digital input of the acquisition board. To synchronize electrophysiological recordings with behavioral data, an LED was positioned within the camera’s field of view in a location not visible to the animal, while behavioral activity was recorded at 30 FPS. The LED was controlled by an Arduino microcontroller, which simultaneously triggered LED illumination and sent a synchronization pulse to the analog input of the Open Ephys acquisition board. A custom Python script was used offline to align video frames containing LED activation with the corresponding analog synchronization signal, enabling precise temporal alignment between electrophysiological and behavioral data. Electrophysiological signals were recorded at a sampling rate of 30 kHz. Animals were habituated to experimenter handling and to the recording apparatus for five days before the start of behavioral procedures.

For LFP analyses, raw traces were visually inspected to identify and exclude channels with excessive noise or unstable signals. Remaining channels were downsampled to 1 kHz and band-pass filtered between 1 and 100 Hz. Object-exploration epochs were manually annotated from video recordings, aligned to the exploration midpoint, and divided into 3-s time windows; corresponding LFP segments were then extracted from the filtered traces for subsequent analyses. LFP power spectral density was estimated using a multitaper method (DPSS tapers) in the 1-100 Hz frequency range. Inter-regional functional coupling was quantified using coherence and Granger causality analyses. Coherence between prefrontal and retrosplenial LFP signals was computed for each exploration epoch across frequencies. Directional interactions were assessed using spectral Granger causality, estimated for each exploration epoch and computed separately for each direction. For band-specific analyses, the theta band was defined as 5-10 Hz, low-gamma as 30-55 Hz, and high-gamma as 55-80 Hz. Absolute LFP power, coherence, and Granger causality values were averaged across these frequency bands for each exploration epoch and then averaged within condition (novel or familiar object exploration).

For single-unit analyses, channels exhibiting excessive noise or unstable recordings were excluded prior to spike sorting. Raw extracellular signals were band-pass filtered between 600 and 6000 Hz, and common-mode noise was attenuated by median subtraction across channels. Recordings were then whitened, and, within each region, the four channels displaying the highest signal variance were selected for spike sorting. Spike detection and automated clustering were then performed using MountainSort5^85^. The resulting spike-sorting output was imported into SpikeInterface for post-processing and quality assessment. Units were initially retained only if they met selected quality criteria (signal-to-noise ratio > 4, ISI violation ratio < 0.01, presence ratio > 0.90, and amplitude cutoff < 0.20). Putative single units were subsequently manually curated in Phy2 by inspecting waveform shape, cluster separation, refractory-period violations, firing stability, and correlogram structure. Only well-isolated units that passed manual curation were retained for subsequent analyses. For peri-event firing analyses, single-unit spike trains were aligned to the video frame corresponding to the onset of exploration for each object. Peri-stimulus time histograms (PSTHs) were computed around each exploration event. A time window from −1 to 0 s relative to exploration onset was defined as the pre-exploration period, whereas the interval from 0 to +1 s was defined as the exploration period.

All electrophysiological analyses were performed in Python using the SpikeInterface, MNE, and SciPy (scipy.signal) libraries.

### Statistical analysis

All data are presented as mean ± SEM. Statistical analyses were performed using GraphPad Prism 8 (GraphPad Software, San Diego, CA), and detailed statistical information for each experiment is provided in the corresponding figure legends. Depending on the experimental design, comparisons were performed using one-way ANOVA, two-way ANOVA, repeated-measures (RM) ANOVA, or two-tailed unpaired t-tests, with Tukey’s and Sidak’s post hoc tests applied after one-way/RM ANOVA and two-way ANOVA, respectively. Nonparametric tests were used when the assumptions for parametric tests were violated. For the brain-wide c-Fos mapping, dimensionality reduction and clustering analyses were conducted in Python using the scikit-learn and UMAP libraries. PCA and Uniform Manifold Approximation and Projection (UMAP) were used for dimensionality reduction and low-dimensional embedding, followed by K-means clustering. Task partial least squares (task-PLS) analysis was performed in Python using a custom script implementing the method described in Franceschini et al. (2023)^8^. For functional network analysis, interregional correlation matrices were computed in Python, and network graphs and topological metrics were obtained using the NetworkX library.

## Acknowledgments

This work was supported by the Ministry of University and Research PNRR project no. (THE) ECS_00000017 and CNR project NutrAge (No. DSB.AD005.225) to N.O.

