## Supplementary Figures for "Distributed neuronal ensembles support episodic-like memory retrieval"

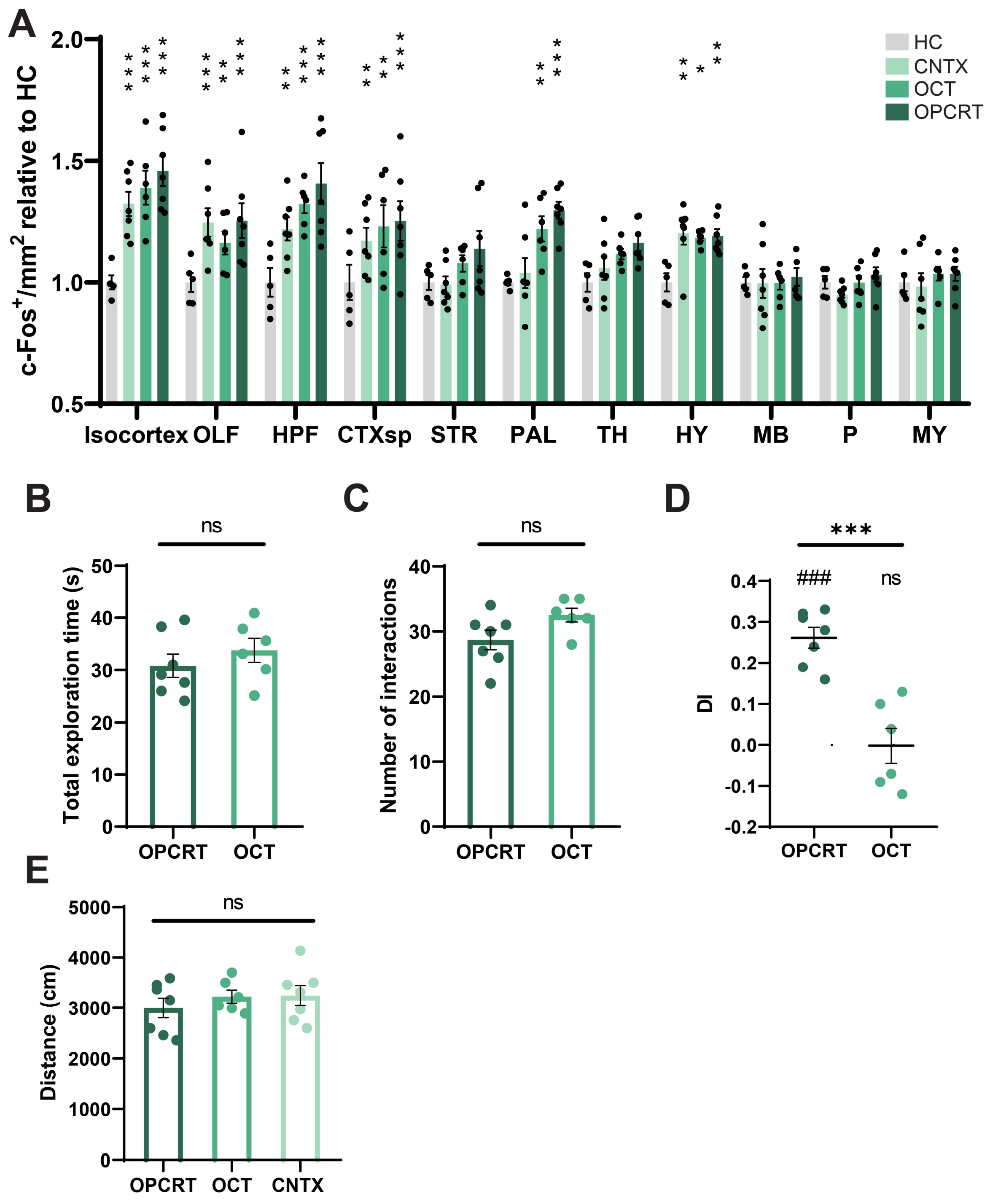


***Figure S1*****:** ***Brain-wide mapping of c-Fos activation during episodic-like memory recall.*** **(A)** c-Fos density normalized to the Home-Cage (HC) control group. Statistical comparisons were performed relative to HC controls; statistical results are reported in Figure 1. **(B)** Total exploration time combined across the two objects for the OPCRT and OCT groups (OPCRT 30.85 ± 2.256, n = 7 vs OCT 33.82 ± 2.301, n = 6, ns p = 0.3786, df = 11, t = 0.9175; unpaired t-test). **(C)** Number of interactions combined across the two objects for the OPCRT and OCT groups (OPCRT 28.71 ± 1.507, n = 7 vs OCT 32.50 ± 1.057, n = 6, ns p = 0.0723, df = 11, t = 1.988; unpaired t-test). **(D)** The OPCRT group showed a significant preference for the object in the novel configuration, as quantified by the Discrimination Index (DI), whereas the OCT group showed no preference between the two identical objects (OPCRT 0.261 ± 0.252, n = 7, *** p < 0.001, df = 6, t = 10.37; OCT -0.001 ± 0.043, n = 6, ns p = 0.9707, df = 5, t = 0.038; one-sample t test. OPCRT vs OCT, *** p < 0.001, df = 11, t = 5.458; unpaired t-test). **(E)** Total distance traveled in the arena, shown for each experimental group (one-way ANOVA with

Sidak's multiple comparisons test: OPCRT 2999 ± 194.1 n = 7 vs OCT 3226 ± 129.6 n = 6, ns p = 0.7804; OPCRT vs CNTX 3249 ± 198 n = 7, ns p = 0.7029; OCT vs CNTX, ns p = 0.9997). Data are presented as mean ± SEM.

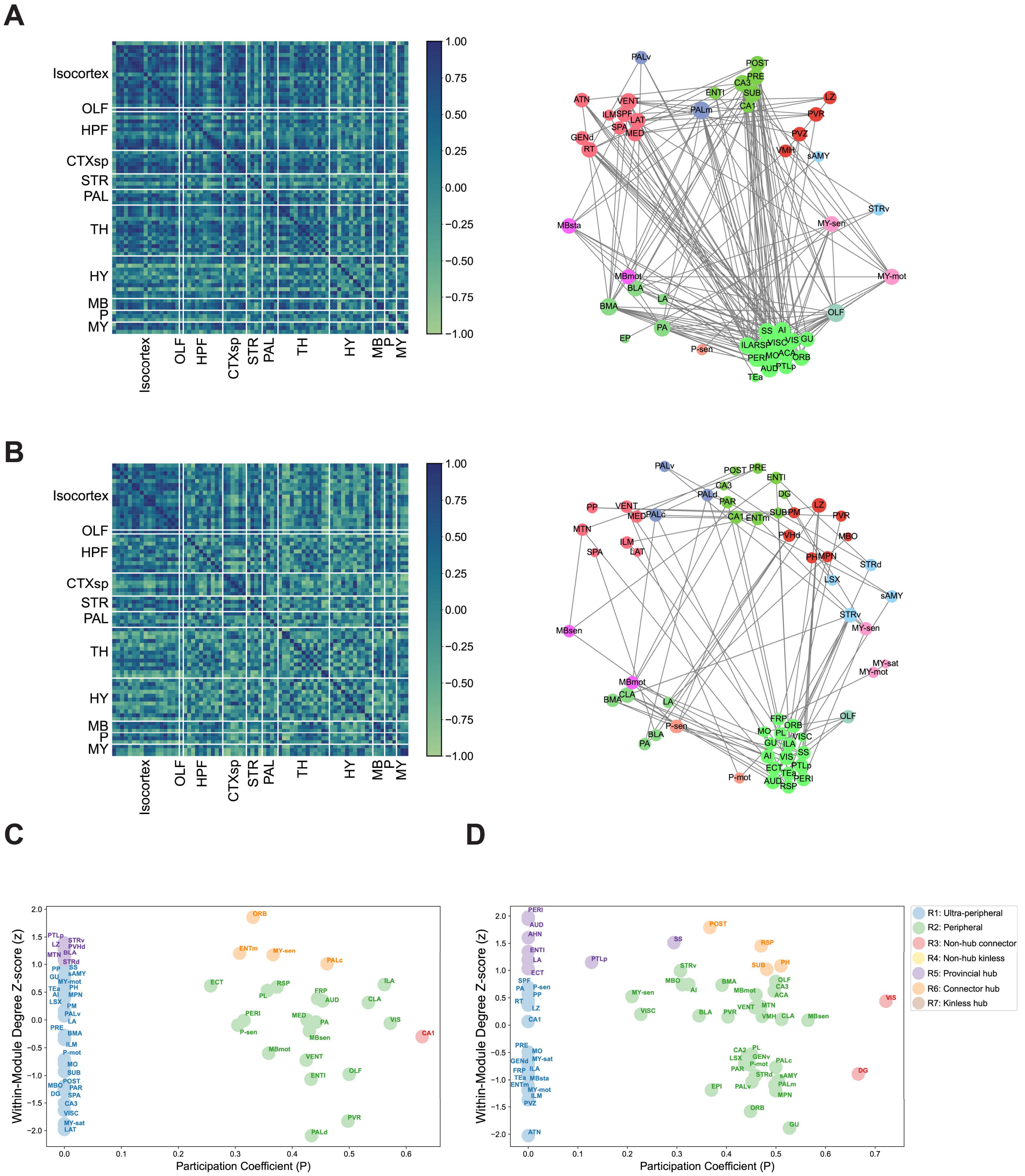

***Figure S2: Topological analysis of functional brain networks.*** Correlation matrices and corresponding network graphs for the CNTX **(A)** and OCT **(B)** groups. Correlation matrices represent pairwise Pearson correlation coefficients computed across brain regions. Functional connectivity networks were generated by retaining only statistically significant correlations ($p < 0.05$) exceeding a strong correlation threshold ($r > 0.8$). Identification of hub regions and topological roles in the CNTX **(C)** and OCT **(D)** functional networks is shown. Nodes were classified according to their within-module degree z-score and participation coefficient, enabling assignment to specific topological roles.

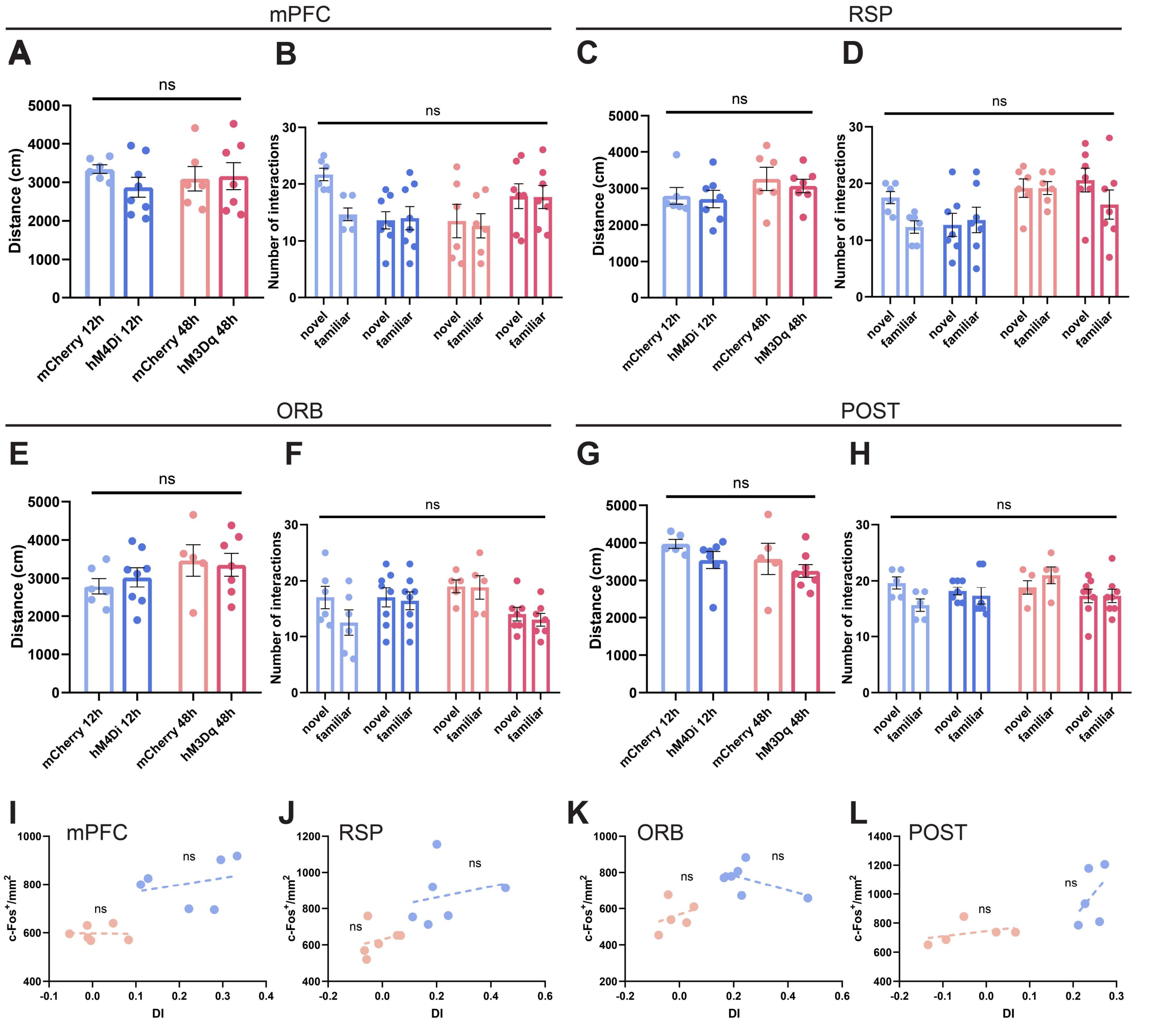


***Figure S3: Chemogenetic manipulation of learning-tagged neuronal populations during OPC memory retrieval.*** Quantification of distance traveled and number of interactions with the objects during the recall phase across the four brain regions analyzed. mPFC: **(A)** total distance traveled in the arena (one-way ANOVA with Sidak's multiple comparisons test: mCherry 12h 3345 ± 112.4 n = 6 vs hM4Di 12h 2874 ± 258.1 n = 8, ns p = 0.8142; mCherry 12h vs mCherry 48h 3094 ± 316.9 n = 6, ns p = 0.9924; hM4Di 12h vs hM3Dq 3159 ± 350.2 n = 7, ns p = 0.9948; mCherry 48h vs hM3Dq 48h, ns p = 0.9999); **(B)** number of interactions (two-way ANOVA with Sidak's multiple comparisons test: mCherry 12h Novel 13.5 ± 2.93 n = 6 vs mCherry 12h Familiar 12.667 ± 2.124 n = 6, ns p = 0.0845; hM4Di 12h Novel 13.625 ± 1.511 n = 8 vs hM4Di 12h Familiar 14 ± 2.044 n = 8, ns p = 0.9998; mCherry 48h Novel 13.5 ± 2.93 n = 6 vs mCherry 48h Familiar 12.667 ± 2.124 n = 6, ns p = 0.9976; hM3Dq 48h Novel 17.857 ± 2.176 n = 7 vs hM3Dq 48h 12h Familiar 17.714 ± 2.032 n = 7, ns p = 0.9999). RSP: **(C)** total distance traveled in the arena (one-way ANOVA with Sidak's multiple comparisons test: mCherry 12h 2799 ± 229.5 n = 6 vs hM4Di 12h 2710 ± 238 n = 7, ns p = 0.9999; mCherry 12h vs mCherry 48h 3262 ± 319.5 n = 6, ns p = 0.7555; hM4Di 12h vs hM3Dq 3068 ± 185.2 n = 7, ns p = 0.8737; mCherry 48h vs hM3Dq 48h, ns p = 0.9944); **(D)** number of interactions (two-way ANOVA with Sidak's multiple comparisons test: mCherry 12h Novel 17.5 ± 1.057 n = 6 vs mCherry 12h Familiar 12.333 ± 1.085 n = 6, ns p = 0.2541; hM4Di 12h Novel 12.714 ± 2.02 n = 7 vs hM4Di 12h Familiar 13.571 ± 2.235 n = 7, ns p = 0.9955; mCherry 48h Novel 19.167 ± 1.621 n = 6 vs mCherry 48h Familiar 19.167 ± 1.138 n = 6, ns p = 0.9999; hM3Dq 48h Novel 20.571 ± 2.091 n = 7 vs hM3Dq 48h 12h Familiar 16.286 ± 2.561 n = 7, ns p = 0.3557). ORB: **(E)** total distance traveled in the arena (one-way ANOVA with Sidak's multiple comparisons test: mCherry 12h 2786 ± 206.1 n = 6 vs hM4Di 12h 3024 ± 251.5 n = 8, ns p = 0.9923; mCherry 12h vs mCherry 48h 3466 ± 410.2 n = 5, ns p = 0.6000; hM4Di 12h vs hM3Dq 3354 ± 299.1 n = 7, ns p = 0.8864; mCherry 48h vs hM3Dq 48h, ns p = 0.9999); **(F)** number of interactions (two-way ANOVA with Sidak's multiple comparisons test: mCherry 12h Novel 17 ± 2.033 n = 6 vs mCherry 12h Familiar 12.500 ± 2.262 n = 6, ns p = 0.2759; hM4Di 12h Novel 17 ± 1.711 n = 8 vs hM4Di 12h Familiar 16.375 ± 1.592 n = 8, ns p = 0.9974; mCherry 48h Novel 19 ± 1.183 n = 5 vs mCherry 48h Familiar 18.8 ± 2.131 n = 5, ns p = 0.9999; hM3Dq 48h Novel 14 ± 1.215 n = 7 vs hM3Dq 48h 12h Familiar 13 ± 1.134 n = 7, ns p = 0.9876). POST: **(G)**

total distance traveled in the arena (one-way ANOVA with Sidak's multiple comparisons test: mCherry 12h 3975 ± 120.2 n = 5 vs hM4Di 12h 3546 ± 226.5 n = 7, ns p = 0.9923; mCherry 12h vs mCherry 48h 3575 ± 412.7 n = 5, ns p = 0.6000; hM4Di 12h vs hM3Dq 3254 ± 168 n = 8, ns p = 0.8864; mCherry 48h vs hM3Dq 48h, ns p = 0.9999); **(H)** number of interactions (two-way ANOVA with Sidak's multiple comparisons test: mCherry 12h Novel 19.6 ± 1.122 n = 5 vs mCherry 12h Familiar 15.6 ± 1.122 n = 5, ns p = 0.1746; hM4Di 12h Novel 18.143 ± 0.705 n = 7 vs hM4Di 12h Familiar 17.286 ± 1.491 n = 7, ns p = 0.9760; mCherry 48h Novel 18.8 ± 1.241 n = 5 vs mCherry 48h Familiar 21 ± 1.483 n = 5, ns p = 0.7103; hM3Dq 48h Novel 17.25 ± 1.221 n = 8 vs hM3Dq 48h 12h Familiar 17.524 ± 1.176 n = 8, ns p = 0.9999). **(I-L)** Relationship between c-Fos density and behavioral performance (DI) across the four brain regions considered (mPFC: mCherry 12h $R^2$ = 0.0747, p = 0.6000; mCherry 48h $R^2$ = 0.0008, p = 0.9556; RSP: mCherry 12h $R^2$ = 0.0474, p = 0.6786; mCherry 48h $R^2$ = 0.0659, p = 0.6232; ORB: mCherry 12h $R^2$ = 0.2897, p = 0.2706; mCherry 48h $R^2$ = 0.1004, p = 0.6035; POST: mCherry 12h $R^2$ = 0.2078, p = 0.4404; mCherry 48h $R^2$ = 0.1601, p = 0.5044). Data are presented as mean ± SEM.

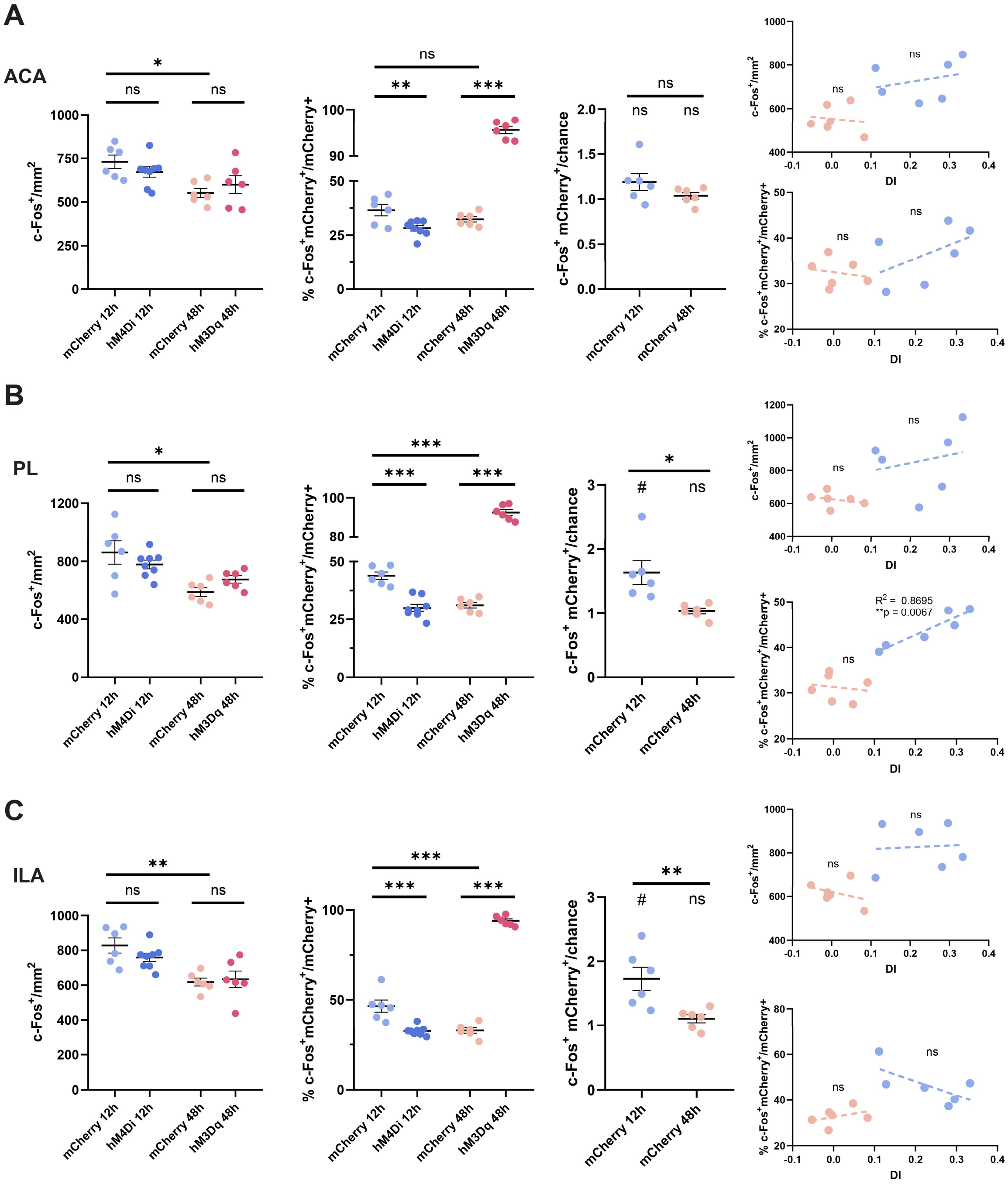


***Figure S4: c-Fos density and ensemble reactivation across selected brain regions during OPC memory retrieval.*** ACA: **(A)** c-Fos+ cell density did not differ following inhibitory or excitatory chemogenetic manipulation, but was significantly higher in the mCherry 12h control group compared to the mCherry 48h control group (one-way ANOVA with Sidak's multiple comparisons test: mCherry 12h 730.7 ± 37.84 n = 6 vs hM4Di 12h 672.5 ± 23.65 n = 8, ns p = 0.8363; mCherry 48h 551.9 ± 26.39 n = 6 vs hM3Dq 48h 599.9 ± 51.14 n = 6, ns p = 0.1345; mCherry 12h vs mCherry 48h, ** p = 0.0185). The proportion of ensemble reactivation was significantly reduced following chemogenetic inhibition and significantly increased following chemogenetic stimulation relative to their respective controls, while no significant difference was observed between the two control groups (one-way ANOVA with Sidak's multiple comparisons test: mCherry 12h 36.54 ± 2.60 n = 6 vs hM4Di 12h 28.28 ± 1.28 n = 8, ** p = 0.0054; mCherry 48h 32.36 ± 1.26 n = 6 vs hM3Dq 48h 95.59 ± 0.79 n = 6, *** p < 0.001; mCherry 12h vs mCherry 48h, ns p = 0.3079). Ensemble reactivation did not differ from chance level in either control group (mCherry 12h 1.189 ± 0.009, n = 6, ns p = 0.0987, df = 5, t = 2.025; mCherry 48h 1.038 ± 0.037, n = 6, ns p = 0.3591, df = 5, t = 1.009; one-sample t test. mCherry 12h vs mCherry 48h, ns p =0.1626, df = 10, t = 1.507; unpaired t-test). No significant correlation was observed between DI and c-Fos density (top right) (mCherry 12h $R^2$ = 0.0811, p = 0.5842; mCherry 48h $R^2$ = 0.0178, p = 0.8010), nor between DI and ensemble reactivation for either control group (bottom right) (mCherry 12h $R^2$ = 0.2637, p = 0.2975; mCherry 48h $R^2$ = 0.039, p = 0.7076). PL: **(B)** c-Fos density did not differ following inhibitory or excitatory chemogenetic manipulation, but was significantly higher in the mCherry 12h control group compared to the mCherry 48h control group (one-way ANOVA with Sidak's multiple comparisons test: mCherry 12h 861.1 ± 80.18 n = 6 vs hM4Di 12h 779 ± 30.02 n = 8, ns p = 0.5566; mCherry 48h 589.5 ± 29.80 n = 6 vs hM3Dq 48h 675.9 ± 25.48 n = 6, ns p = 0.5698; mCherry 12h vs mCherry 48h, ** p = 0.0025). Ensemble reactivation was significantly reduced following chemogenetic inhibition and significantly increased following chemogenetic stimulation relative to their respective controls. Ensemble reactivation differed significantly between the two control groups, being higher in mCherry 12h than in mCherry 48h (one-way ANOVA with Sidak's multiple comparisons test: mCherry 12h 43.88 ± 1.6 n = 6 vs hM4Di 12h 30.04 ± 1.28 n = 8, *** p < 0.001; mCherry 48h 31.20 ± 1.21 n = 6 vs hM3Dq 48h 92.41 ± 1.57 n = 6, *** p < 0.001; mCherry 12h vs mCherry 48h, *** p < 0.001). Reactivation in the mCherry 12h group was also significantly higher than both chance level and the mCherry 48h group (mCherry 12h 1.264 ± 0.185, n = 6, * p = 0.0186, df = 5, t = 3.430; mCherry 48h 1.166 ± 0.045, n = 6, ns p = 0.4912, df = 5, t = 0.7424; one-sample t test. mCherry 12h vs mCherry 48h, * p =0.0103, df = 10, t = 3.151; unpaired t-test). No significant correlation was observed between DI and c-Fos density in either control group (mCherry 12h $R^2$ = 0.0530, p = 0.6606; mCherry 48h $R^2$ = 0.0828, p = 0.3614); however, a significant positive correlation was observed between ensemble reactivation and DI in the mCherry 12h group (mCherry 12h $R^2$ = 0.8695, p = 0.0067; mCherry 48h $R^2$ = 0.0252, p = 0.7636). ILA: **(C)** c-Fos density did not differ following inhibitory or excitatory chemogenetic manipulation, but was significantly higher in the mCherry 12h control group compared to the mCherry 48h control group (one-way ANOVA with Sidak's multiple comparisons test: mCherry 12h 827.9 ± 43.45 n = 6 vs hM4Di 12h 759.1 ± 24.12 n = 8, ns p = 0.4905; mCherry 48h 617.6 ± 22.55 n = 6 vs hM3Dq 48h 633.5 ± 47.63 n = 6, ns p = 0.9893; mCherry 12h vs mCherry 48h, ** p = 0.0024). The proportion of ensemble reactivation was significantly reduced following chemogenetic inhibition and significantly increased following chemogenetic stimulation relative to their respective controls, with greater reactivation in mCherry 12h compared to mCherry 48h (one-way ANOVA with Sidak's multiple comparisons test: mCherry 12h 46.48 ± 3.38 n = 6 vs hM4Di 12h 32.56 ± 0.91 n = 8, *** p < 0.001; mCherry 48h 32.78 ± 1.58 n = 6 vs hM3Dq 48h 93.92 ± 1.11 n = 6, *** p < 0.001; mCherry 12h vs mCherry 48h, *** p < 0.001). Reactivation exceeded chance levels in the mCherry 12h group but not in the mCherry 48h group, and the two control groups differed significantly (mCherry 12h 1.728 ± 0.181, n = 6, * p = 0.0103, df = 5, t = 4.006; mCherry 48h 1.106 ± 0.063, n = 6, ns p = 0.1577, df = 5, t = 1.661; one-sample t test. mCherry 12h vs mCherry 48h, ** p =0.0090, df = 10, t = 3.231; unpaired t-test). In contrast to the PL, no significant association was detected between DI and either c-Fos density (mCherry 12h $R^2$ = 0.004, p = 0.8979; mCherry 48h $R^2$ = 0.1242, p = 0.4933).) or ensemble reactivation (mCherry 12h $R^2$ = 0.4589, p = 0.1393; mCherry 48h $R^2$ = 0.1414, p = 0.4626).) in this subregion. Data are presented as mean ± SEM.
